# Integrated Stress Response Deregulation underlies Vanishing White Matter and is a target for therapy

**DOI:** 10.1101/460840

**Authors:** Truus E.M. Abbink, Lisanne E. Wisse, Ermelinda Jaku, Michiel J. Thiecke, Daniel Voltolini-González, Hein Fritsen, Sander Bobeldijk, Timo J. ter Braak, Emiel Polder, Nienke L. Postma, Marianna Bugiani, Eduard A. Struijs, Mark Verheijen, Nina Straat, Sophie van der Sluis, Adri A.M. Thomas, Douwe Molenaar, Marjo S. van der Knaap

**Author notes:** To whom correspondence should be addressed: Dr. Truus E.M. Abbink, De Boelelaan 1085 WN-B643, 1081 HV, Amsterdam, the Netherlands. current affiliation, Nuclear Dynamics Programme, The Babraham Institute, Babraham Research Campus, Cambridge, UK. current affiliation, Sylics, Amsterdam, the Netherlands (Nina Straat).

## Abstract

Vanishing white matter (VWM) is a fatal, stress-sensitive leukodystrophy that mainly affects children and is currently without treatment. VWM is caused by recessive mutations in eukaryotic initiation factor 2B (eIF2B) that is crucial for initiation of mRNA translation and its regulation under stress conditions. Mice with bi-allelic missense mutations in eIF2B recapitulate human VWM. VWM pathomechanisms are unclear. Using polysomal profiling to screen for mRNAs with altered translation we observed most prominent changes in expression of integrated stress response (ISR) mRNAs in brains of mutant compared to wild-type mice; expression levels correlated with disease severity. We substantiated these findings in VWM patients’ brains. ISRIB, an ISR inhibitor, normalized expression of mRNA markers, ameliorated white matter pathology and improved motor skills in VWM mice, thus showing that the ISR is central in VWM pathomechanisms and a viable target for therapy.

**One Sentence Summary:** ISRIB ameliorates ISR deregulation and clinical signs in VWM mice

## Introduction

Vanishing white matter (VWM) is a leukodystrophy, a genetic disorder of the white matter of the central nervous system, mainly affecting young children (*1*). Patients with VWM display chronic neurological deterioration with additional episodes of rapid and severe decline provoked by stresses, especially febrile infections (*1, 2*). Neuropathology of post-mortem brain shows white matter rarefaction and cystic degeneration with feeble astrogliosis, profound lack of myelin, and immature astrocytes and oligodendrocytes (*3*). There is no cure for VWM and patients die prematurely. Recessive mutations in any of the 5 genes encoding the subunits of eukaryotic translation factor 2B (eIF2B) cause VWM (*4, 5*). Mutations reduce the activity of eIF2B (*6-8*). eIF2B is conditional for mRNA translation initiation *per se* and essential for regulating protein synthesis rates and orchestrating the integrated stress response (ISR, Fig.1a) (*9*). VWM mouse models with homozygous (ho) missense mutations in eIF2Bδ (Arg484Trp) or eIF2Bε (Arg191His) subunits are representative of the human disease (*2b4*^*ho*^ and *2b5*^*ho*^ mice) (*10*). These mutations correspond to the eIF2Bδ Arg483Trp and eIF2Bε Arg195His mutation in patients. Each mutation reduces eIF2B activity in biochemical assays and causes a severe VWM phenotype in patients (*11, 12*). Crossbreeding generates *2b4*^*he*^*2b5*^*ho*^ and *2b4*^*ho*^*2b5*^*he*^ and *2b4*^*ho*^*2b5*^*ho*^ (he, heterozygous) mice. Together, the mouse models reproduce the clinical spectrum of human VWM in the following order of increasing severity: *2b4*^*ho*^, *2b5*^*ho*^, *2b4*^*he*^*2b5*^*ho*^/*2b4*^*ho*^*2b5*^*he*^ and *2b4*^*ho*^*2b5*^*ho*^, as reflected by earlier disease onset, more severe motor dysfunction, lower body weight, shorter life span, and severer brain pathology (*10*). Using these mice, we showed that astrocytes are the primarily affected cell type (*10*).

Considering the housekeeping function of eIF2B in all cells, it is remarkable that a defect in eIF2B predominantly affects astrocytes. The underlying mechanism is unclear. We hypothesize that translation of specific mRNAs in brain is altered by eIF2B mutations, primarily impacting astrocytes. In this study we performed polysomal profiling to identify and quantify polysome-associated mRNAs and found differential expression of ATF4-regulated mRNAs in VWM mouse brain, indicative of constitutively enhanced expression of ATF4 and its transcriptome by reduced eIF2B activity. These changes were already seen at an early disease stage with minimal signs of white matter damage. eIF2α phosphorylation was reduced below baseline, indicative of a constitutively activated feedback mechanism, clearly unable to fully compensate for the reduced eIF2B activity. Together the findings revealed a deregulated ISR homeostasis in VWM. The expression level of the ATF4-regulated transcriptome correlated positively with disease development and severity. Immunohistochemistry of VWM mouse brain revealed expression of ATF4 and its target 4EBP1 (*13, 14*) only in astrocytes. We substantiated these findings in VWM patients’ brains. We used ISRIB, an ISR inhibitor (*15-18*), in VWM mice to modulate the deregulated ISR and study the impact of this modulation on the disease. ISRIB stabilizes the eIF2B complex by direct binding (*16, 19, 20*). As a consequence, ISRIB enhances eIF2B activity, which reduces the expression of ATF4 and impacts expression of the ATF4-regulated transcriptome (*16, 18, 19*). The compound has been successful in treatment of a prion-infected mouse model for neurodegeneration (*15*). We revealed that ISRIB normalizes ATF4 expression, astrocyte morphology, brain pathology and motor skills of VWM mice.

## Results

### Polysomal profiling reveals 33 mRNAs with altered translation in brains of 2b5^ho^ versus WT mice

We performed polysomal profiling with forebrains of 4-month-old wild-type (WT) and *2b5*^*ho*^ mice, shortly before onset of clinical signs in *2b5*^*ho*^ mice (*10*). We reasoned that at this age eIF2Bε Arg191His-regulated changes in mRNA translation would be detectable, while general changes in mRNA translation in response to tissue damage would be minimal. The polysomal profiles generated from WT and *2b5*^*ho*^ samples were similar, showing that bulk mRNA translation was not grossly affected by the mutations (Fig. S1). The relative distribution between monosome and polysome fractions was determined for each mRNA per genotype. Differential distribution between the WT and *2b5*^*ho*^ samples was found for 33 mRNAs (Table S1), indicative of altered translation rates in *2b5*^*ho*^ brain; the majority were shifted towards the polysome fraction in *2b5*^*ho*^ mouse forebrain samples indicative of increased translation efficiencies. Strikingly, 17 out of the 33 differentially translated mRNAs are regulated by transcription factor ATF4 (Fig. 1a) (*13*), indicating increased transcription.

**Fig. 1.**
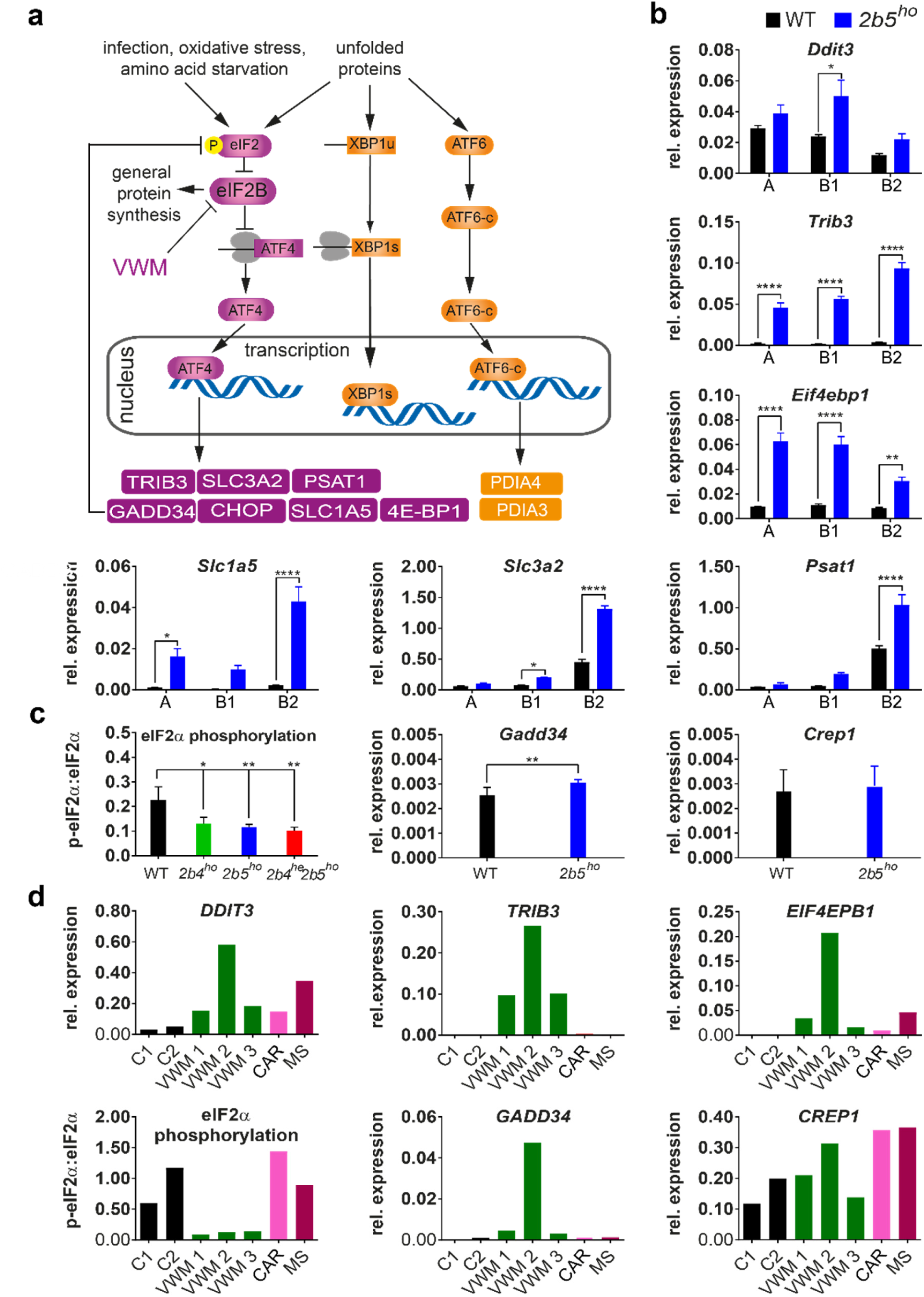
Polysomal profiling of *2b5*^*ho*^ mouse brain identifies ISR deregulation in mouse VWM and human VWM brain. (**a**) Summary of ISR- and UPR-regulated transcription included to clarify the link between the ATF4, ATF6-c and XBP1s transcription factors and mRNA targets investigated in the current study. (**b**) ATF4-regulated mRNA levels in gradient fractions from forebrain lysates from 4-month-old WT and *2b5*^*ho*^ mice were measured to visualize the mRNA distribution in monosome fraction (A), polysome fractions (B1, less than 5 ribosomes per mRNA; B2, 5 or more ribosomes per mRNA). Graphs show average ± sd, n=3 (*Akt*, qPCR reference). Statistical significance was determined by two-way ANOVA with Sidak’s correction; *p<0.05, **p<0.01, ****p<0.0001). (**c**) eIF2α phosphorylation, *Gadd34* and *Crep1* mRNA expression levels were measured in cerebellar tissue from WT and VWM mice, as indicated. Graphs show average ± sd, n=3 for eIF2α phosphorylation and n=6 for mRNA expression (*Gapdh+Akt*, qPCR reference). Statistical differences in eIF2α phosphorylation were determined using a one-way ANOVA followed by a Dunnet’s correction. Differences between WT and *2b5*^*ho*^ qPCR data were assessed with student T-test, *p<0.05, **p<0.01. (**d**) *DDIT3, TRIB3, EIF4EBP1* mRNA levels, eIF2α phosphorylation, *GADD34* and *CREP1* mRNA levels were quantified in postmortem frontal white matter tissue from VWM patients and controls (negative controls without brain pathology, C1, C2 and disease controls CAR and MS; CAR, CARASAL; MS, multiple sclerosis). Expression differences among VWM patients inversely correlate with postmortem delay time (*AKT+GAPDH*, qPCR reference, n=1). Details on control and patients’ tissue are listed in Supplementary File 4.

### Overrepresentation analysis of mRNAs with differential polysome association shows altered ATF4-driven transcription in 2b5^ho^ mouse and VWM patient brain

We evaluated the microarray findings for a number of the candidates with qPCR (Fig. 1b, Fig. S2). The majority of the mRNAs with altered translation rates accumulated at increased levels in monosome and polysome fractions from brain of *2b5*^*ho*^ mice, suggesting that these mRNAs were also regulated at the transcript level. We focused on differences in mRNA levels within the polysome fractions, without taking into account their distribution between the monosome and polysome fractions. We performed overrepresentation analysis with the eIF2Bε^Arg191His^-regulated polysome-associated mRNAs, We identified 176 mRNAs (detected with 234 probes) with an altered level of polysome association in forebrain of *2b5*^*ho*^ compared to WT mice (115 up and 61 down, Suppl. File 2), indicating altered synthesis of the proteins they encode. Overrepresentation analysis (Fig. 2a) highlighted changes in amino acid transport, serine biosynthesis and glycine metabolism. Out of the 176 mRNAs 52 are regulated by ATF4 (*13*), including *Ddit3*, the mRNA that encodes the transcription factor CHOP (Data file S3). ATF4-regulated mRNA’s were enriched 5-fold compared to the mRNAs, for which polysome-association was similar in WT and *2b5*^*ho*^ forebrain (p<0.001; Data file S3). We verified findings from mice in humans (Fig. 1d); expression of *DDIT3, TRIB3* and *EIF4EBP1* mRNA was increased in VWM patients’ brain. Microarray analyses of total RNA from postmortem white matter tissue confirmed a 2-fold enrichment of ATF4-regulated mRNAs in differentially expressed mRNAs in VWM patients compared to controls (p<0.0001; File S3).

**Fig. 2.**
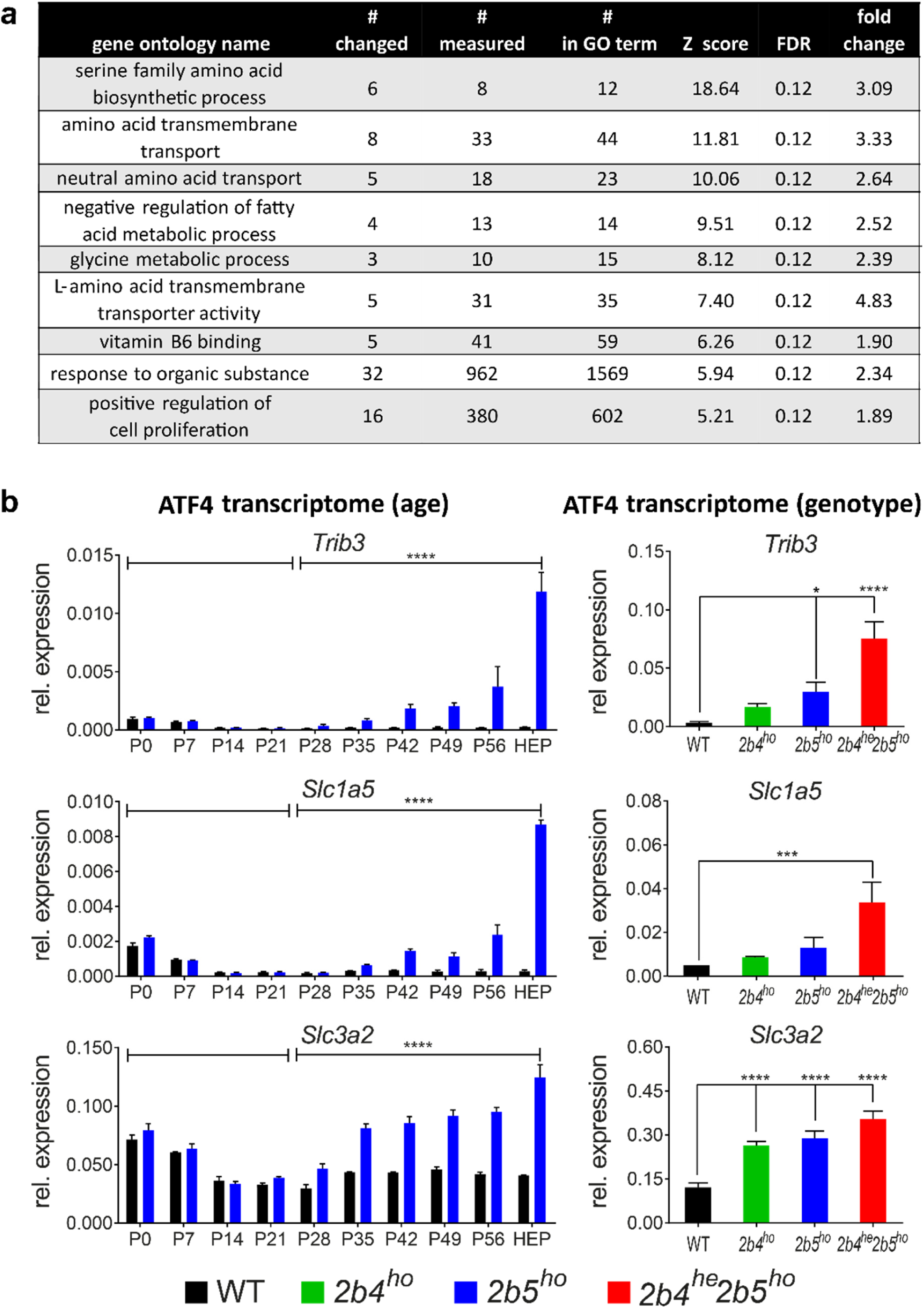
Follow-up analyses show affected cellular functions and ISR deregulation correlating with disease development. (**a**) Overrepresentation analysis of regulated polysome-associated mRNAs in *2b5*^*ho*^ mouse forebrain against the gene ontology (GO) database. # changed, number of mRNAs in GO term differentially associated with *2b5*^*ho*^ polysomes; # measured, number of mRNAs in GO term detected in WT and *2b5*^*ho*^ polysomes; # in ontology, total gene number in GO term; results with a Z score >2 and FDR of < 0.15 were considered significant; fold change, average of differentially expressed genes within the dataset for the particular GO term. (**b**) Expression of the ATF4-regulated transcriptome in VWM mouse brain correlates positively with disease development. mRNA levels were determined at indicated ages (left hand side) or in indicated genotypes (right hand side). Sagittally sliced brain halves of WT and *2b5*^*ho*^ mice were analyzed at indicated ages (*Gapdh*, qPCR reference; P, postnatal day; HEP, humane end point). Graphs show average ± sd, n=2. Statistical analysis on the differences at groups of different ages was done for each mRNA using a two-way ANOVA (genotype*day interaction all days) followed by 2 two-way ANOVAs for each mRNA (one comparing data of <P21 with ≥P21 and one comparing data of <P28 with ≥P28 for genotype*day interaction) using SPSS. Cerebella of indicated mouse genotypes were analyzed at 4-month-old age (*Gapdh*+*Akt*, qPCR reference). Graphs show average ± sd, n=3 (n=4 for *2b4*^*ho*^). Statistical differences were determined using a one-way ANOVA followed by Tukey’s correction. **p<0.01, ***P<0.001, ****p<0.0001.

### Expression of the ATF4 transcriptome is accompanied by reduced eIF2α phosphorylation

We wondered whether the ATF4 transcriptome expression is enhanced in VWM brain due to ISR activation via eIF2α phosphorylation. In contrast with such hypothesis, we found that eIF2α phosphorylation in brain was significantly reduced in VWM compared to the baseline found in WT mice (Fig. 1c, Fig. S3). Dephosphorylation of eIF2α is mediated by protein phosphatase 1, using either CREP1 or GADD34 as co-factor (*21*). GADD34 expression is induced during the ISR in an ATF4- and CHOP-dependent manner; GADD34 is essential for the negative feedback of the ISR and restores protein synthesis (Fig. 1a), while CREP1 functions as a co-factor also in unstressed cells (*13, 22-24*). *Crep1* mRNA expression was similar, but *Gadd34* mRNA expression was increased in *2b5*^*ho*^ brain compared to WT brain (Fig. 1c), in line with its regulation by ATF4 and CHOP (*25*) (Fig. 1a). In VWM patients’ white matter tissue eIF2α phosphorylation was significantly decreased compared to control tissue (Fig. 1d). In line with this observation, *GADD34* but not *CREP1* mRNA levels were increased in VWM patient brain tissue compared to control tissue. The decreased level of eIF2α phosphorylation can be directly explained by increased expression of GADD34, resulting from the increased level of *Gadd34* mRNA we found. Western blot analysis for GADD34 protein was not successful.

The ISR constitutes part of the unfolded protein response (UPR), which is activated by endoplasmic reticulum (ER) stress (Fig. 1a). UPR activation in patients’ white matter was previously reported (*26, 27*). Therefore, we investigated activation of UPR branches regulated by ATF6 or IRE1α in *2b5*^*ho*^ mouse brain. We measured specific markers for ATF6-driven transcription (*Pdia4* mRNA) and IRE1α activation (*Xbp1* mRNA splicing) (Fig. S4) (*28-30*). qPCR analyses did not show increased expression of the *Pdia4* mRNA nor *Xbp1* mRNA splicing, arguing against recent or ongoing activation of the UPR in *2b5*^*ho*^ brain. The previously reported expression of UPR markers in patient brains was determined in end-stage disease (*26, 27, 31*) and could be related to that stage. We measured *Pdia4* mRNA expression and *Xbp1* mRNA splicing in mouse brain at the humane end point. Still, we did not observe an increased expression of the UPR markers compared to the WT brain (Fig. S4). We further verified findings in human tissue and found comparable expression of *PDIA3* and *XBP1* splicing in white matter tissue from controls and VWM patients (Fig. S4). Microarray analyses of total RNA from postmortem white matter tissue did not show enrichment of ATF6- and/or XBP1-regulated mRNAs either in differentially expressed mRNAs in VWM patients compared to controls (p=0.4507; Data file S3). These findings indicate that eIF2B mutations do not chronically activate ER stress but alter the ISR homeostasis leading to a deregulated state.

### Expression of the ATF4 transcriptome in 2b5^ho^ mouse brain correlates with disease development and severity

We assessed expression levels for several mRNA candidates in mouse brain at ages ranging from postnatal day 0 (P0) to the humane end point (Fig. 2b). From P0 to P14 none of the tested candidates differed in expression level between WT and *2b5*^*ho*^ brain (Suppl. Data file 5). From P28 onwards the expression of ATF4-regulated mRNAs changed progressively in *2b5*^*ho*^ but not in WT brain (p<0.0001; Fig. 2b; Suppl. Data file 5). The maximal expression differences were measured at the humane end point.

We also measured the expression of several ATF4-regulated mRNAs in brains of *2b4*^*ho*^, *2b5*^*ho*^ and *2b4*^*he*^*2b5*^*ho*^ mice, displaying a severity range from mild to severe (Fig. 2b) (*10*). The expression of several mRNA markers correlated positively with disease severity in the mouse models (Data file S5). We did not detect ISR mRNA marker expression in brain tissue of an unrelated neurodegenerative mouse model (Fig. S5) (*32, 33*). In addition, brain tissue samples from patients with multiple sclerosis (*34*) or cathepsin A-related arteriopathy with strokes and leukoencephalopathy (CARASAL) (*35*) did not show consistently altered expression of ISR markers (Fig. 1d, Fig. S4), indicating that ISR deregulation is not a general event in white matter disease.

### ATF4 and 4E-BP1 expression in white and gray matter astrocytes

To identify which specific cells or brain areas display an activated ATF4 transcriptome we performed immunohistochemistry (IHC) for ATF4 as well as ATF4-regulated 4E-BP1 on WT and VWM mouse brain (Fig. 3). The tested antibodies for CHOP, TRIB3 and SLC3A2 did not yield specific signals and were not used. IHC for ATF4 on WT and *2b5*^*ho*^ brain showed nuclear ATF4 immunoreactivity in cerebellum, corpus callosum and cortex in *2b5*^*ho*^ brain only (Fig. 3). Western blot analyses on brain nuclei confirmed ATF4 specificity of the antibody (Fig. S6). The morphology of the ATF4-positive nuclei, based on size, degree of chromatin compaction and presence of nucleoli, indicates that the positive cells belong to the macroglial lineage. IHC for 4EBP1 on WT, *2b4*^*ho*^, *2b5*^*ho*^ and *2b4*^*he*^*2b5*^*ho*^ showed 4E-BP1 immunoreactivity in both white and grey matter cells of cerebellum, corpus callosum and cortex for the VWM mouse brains but not for WT mouse brain (Fig. 3). The latter is not a surprise as 4E-BP1 is not expressed in WT brain, which expresses 4E-BP2 (*36*). Astrocytes in brain sections from 1 month-old *2b5*^*ho*^ mice were already positive for 4E-BP1 (data not shown). The morphology of the 4E-BP1-positive cells corresponded to white and grey matter astrocytes, including Bergmann glia. To confirm the astrocytic expression of both ATF4 and 4E-BP1, double-stainings were performed with antibodies against ATF4 or 4E-BP1 combined with an antibody against astrocyte marker GFAP in *2b4*^*he*^*2b5*^*ho*^ or *2b5*^*ho*^ mice, respectively. The stainings identified ATF4-GFAP and 4E-BP1-GFAP double-positive cells in brain sections of VWM mice but not of WT mice (Fig. S7). No cells with the morphology of oligodendrocytes and positive for 4E-BP1 were detected in *2b4*^*ho*^, *2b5*^*ho*^ and *2b4*^*he*^*2b5*^*ho*^ brain, even at the humane end point (Fig. 3). Double-stainings with antibodies against 4E-BP1 and Olig2, as marker for oligodendrocytes (*37*), did not show double-positive cells in *2b5*^*ho*^ mouse brain (Fig. S7). Double-stainings with antibodies against Olig2 and ATF4 to investigate if ATF4 is expressed in oligodendrocytes are technically challenging. The Olig2 signal is much higher than the ATF4 signal and both signals localize in the nuclei, potentially causing false-negative findings. Immunofluorescence would overcome this problem, but ATF4 was not detected in this type of assay in our hands. Co-stainings for Olig2 and 4E-BP1 were thus favored, as the signal for 4E-BP1 localizes to the cytoplasm and not to the nucleus. 4E-BP1-positive cells with the morphology of astrocytes were also detected in the frontal white matter, cerebellum, and cortex of post-mortem brain tissue from VWM patients (Fig. 3). 4E-BP1-positive cells with the morphology of oligodendrocytes or neurons were not observed in VWM patient brain tissue. 4E-BP1-positive astrocytes, oligodendrocytes or neurons were not observed in brains from MS or CARASAL patients (data not shown). Staining with the ATF4 antibody did not give a specific signal in human brain and prevented determination of the localization of ATF4 protein in human VWM brain sections.

**Fig. 3.**
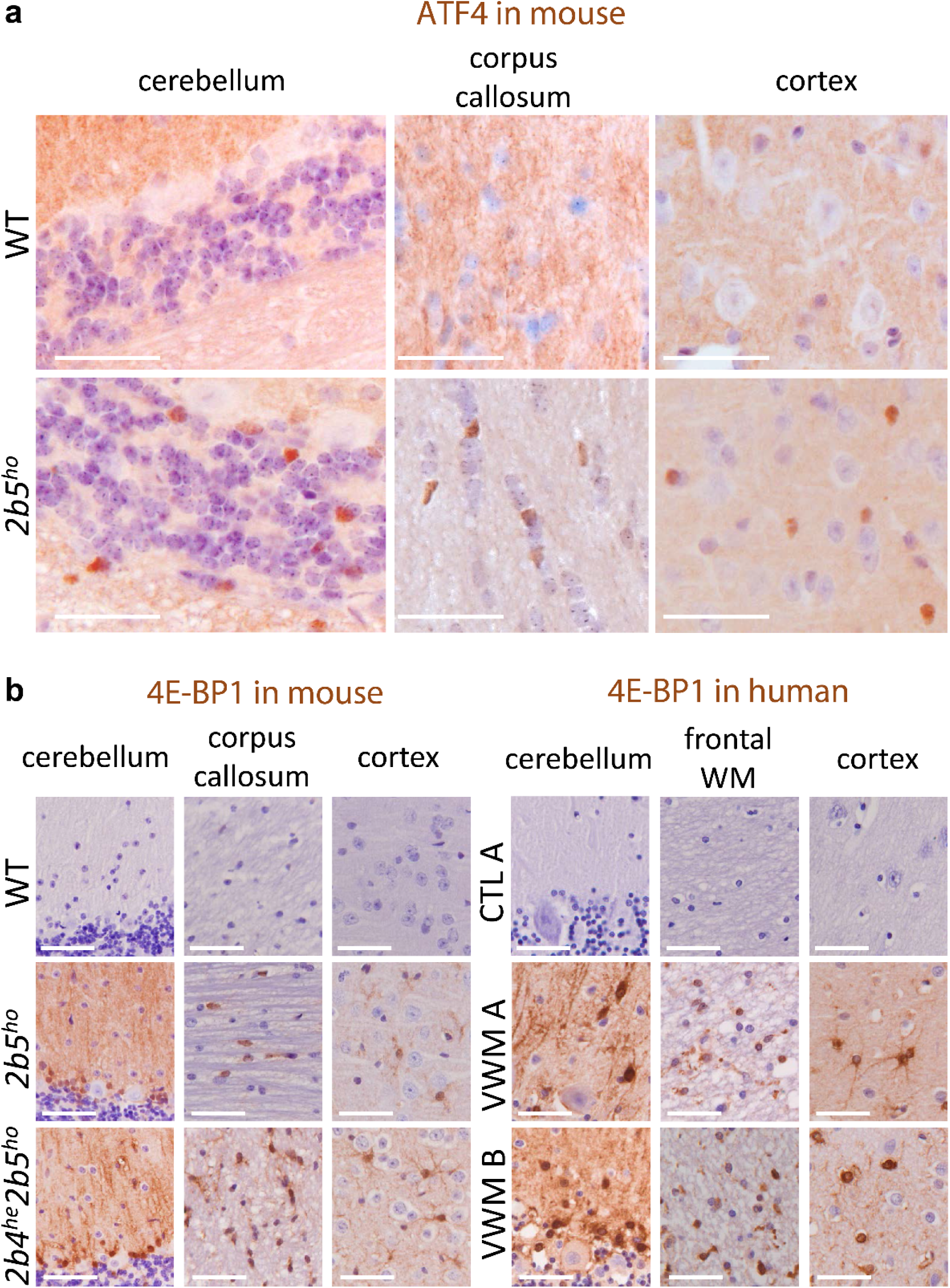
White and grey matter astrocytes in VWM mice and VWM patients are immunoreactive for ATF4-regulated 4E-BP1. (**a**) ATF4 immunoreactive nuclei (brown) are detected in white and grey matter macroglia of VWM mice. Sections from brain tissue from 4-month-old WT and *2b5*^*ho*^ mice were stained with antibodies against ATF4. Findings in cerebellum, corpus callosum and cortex are indicated. Staining for ATF4 on human brain sections was not successful. (**b**) White and grey matter astrocytes in VWM mouse and VWM human brain in show 4E-BP1 immunoreactivity (brown). Brain sections from 4-month-old mice (WT, *2b5*^*ho*^ and *2b4* ^*he*^*2b5*^*ho*^), human control (C1) or patients (VWM 1 and VWM 2) were stained with antibodies against 4E-BP1. Details on human brain tissue are listed in Supplementary File 4. Findings in cerebellum, corpus callosum (mouse) or frontal white matter (WM, human) and cortex are shown. Purple stain indicates nuclei. White bar, 0.05 mm.

### ISRIB injections differentially normalize clinical signs of VWM mice

We postulate that expression of the ATF4 transcriptome in astrocytes underlies their dysfunction. We targeted the ISR with ISRIB (*15-18, 38*) to reduce the expression of the ATF4 transcriptome and assess its role in pathogenesis. We developed a vehicle to overcome ISRIB’s insolubility, which hinders bio-availability in mice (*15*). ISRIB was administered daily to male WT, *2b5*^*ho*^ and *2b4*^*ho*^*2b5*^*he*^ mice by intraperitoneal (i.p.) injections. The age of the mice at the start of the injection scheme ranged from 6 to 8 weeks. We selected the *2b5*^*ho*^ and *2b4*^*ho*^*2b5*^*he*^ genotypes to determine ISRIB’s effect on the individual mutations: the severely affected *2b4*^*ho*^*2b5*^*he*^ mice were selected over the mildly affected *2b4*^*ho*^ mice, as in *2b4*^*ho*^ mice daily injections would be required for many months (approximately 16-18). Injecting ISRIB into *2b5*^*ho*^ and *2b4*^*ho*^*2b5*^*he*^ mice allowed assessing ISRIB’s effects on a moderate and severe variant of VWM disease.

ISRIB increased VWM but not WT mouse body weight compared to saline-injected genotype-matched animals (Fig. 4a). The weight gain was highest in *2b4*^*ho*^*2b5*^*he*^ mice and was detected after a few days of injections (Fig. 4a). We continued daily injections, and neuroscores were taken to determine signs of neurological deterioration in VWM mice^32^. Neurological deterioration was observed in placebo-treated *2b5*^*ho*^ and *2b4*^*ho*^*2b5*^*he*^ mice after 18-20 or 11-12 weeks of injections (Fig. 4b). ISRIB-treated *2b5*^*ho*^ mice showed signs of mild neurological deterioration when reaching an age of 40 weeks (32-34 weeks of injection, Fig. 4b). ISRIB-treated *2b4*^*ho*^*2b5*^*he*^ mice did not show neurological signs for the duration of the experiment (Fig. 4b). At the end of the experiment, ISRIB’s effects were measured in several motor tests. ISRIB improved performance of VWM mice in Balance Beam tests, in which the primary outcome, number of slips, was significantly decreased compared to placebo-treated VWM mice (Fig. 4c). Static and dynamic gait parameters were investigated with the catwalk. Most gait parameters in placebo-treated *2b4*^*ho*^*2b5*^*he*^ mice were more affected than placebo-treated *2b5*^*ho*^ mice (Fig. 4d; Fig. S8; Data file S5). ISRIB improved all these parameters in *2b4*^*ho*^*2b5*^*he*^ mice reaching WT levels (Fig. 4d; Fig. S8; Data file S5). ISRIB effects on gait parameters in *2b5*^*ho*^ mice were less consistent and less pronounced than in *2b4*^*ho*^*2b5*^*he*^ mice (Fig. 4d; Fig. S8; Data file S5). ISRIB did not affect any parameters in WT mice (Fig. 4d; Fig. S8; Data file S5).

**Fig. 4.**
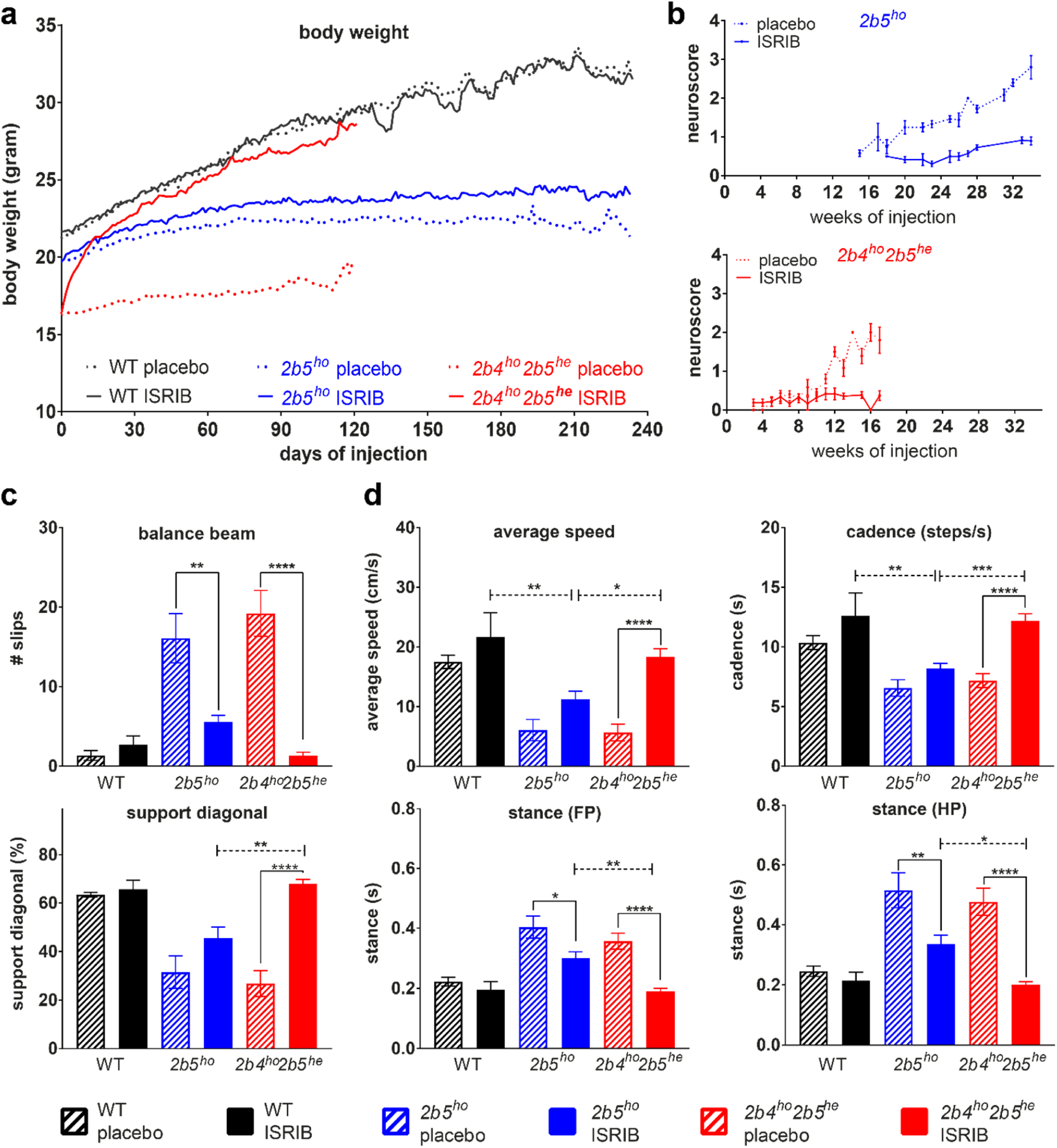
ISRIB ameliorates clinical signs in two VWM mouse models, most effectively in *2b4*^*ho*^*2b5*^*he*^ mice. Mice were injected daily with vehicle or 1mg/kg ISRIB. (**a**, **b**, **c**, **d**) Graphs show phenotypic measures of placebo- and ISRIB-treated WT (n=6 per condition) and VWM mice (*2b5*^*ho*^ and *2b4*^*ho*^*2b5*^*he*^ n=14 per condition or as indicated): average body weight of WT and VWM mice (**a**), average neuroscore in VWM mice (**b**), average number of slips on balance beam (*2b5*^*ho*^ n=11 for placebo, n=13 for ISRIB and *2b4*^*ho*^*2b5*^*he*^ n=10 for placebo and n=13 for ISRIB, **c**), average measures of selected CatWalk parameters (*2b5*^*ho*^ n=9 for placebo, n=13 for ISRIB and *2b4*^*ho*^*2b5*^*he*^ n=12 for placebo and n=14 for ISRIB, **d**). Error bars indicate SD (graph b), SEM (graphs in c, d) or are left out (graph in a) to allow visualization of the mean values (SEM is shown in Suppl. Data File 5). Raw data of all CatWalk parameters are given in Supplementary File 5. Statistical analysis investigating the ISRIB differences in WT, *2b5*^*ho*^ and *2b4*^*ho*^*2b5*^*he*^ was performed with a two-way ANOVA with Tukey’s correction (Suppl. Data File 5). *, p<0.05, **p<0.01, ***P<0.001, ****p<0.0001.

### ISRIB impacts expression of ISR mRNA markers and improves histopathology

Expression of several ATF4-regulated mRNAs was measured in cerebella, collected from placebo- and ISRIB-injected mice at the end of the experiment. ISRIB reduced expression of *Atf4* and ATF4-regulated mRNAs in the VWM mice and most consistently in *2b4*^*ho*^*2b5*^*he*^ mice (Fig. 5a). ISRIB did not affect the phosphorylation of eIF2α in *2b5*^*ho*^ and *2b4*^*ho*^*2b5*^*he*^ mice, despite reduced *Gadd34* mRNA levels (Fig. 5a; Fig. S9). ISRIB reduced expression of the *Eif4ebp1* mRNA and 4E-BP1 protein in ISRIB-injected *2b4*^*ho*^*2b5*^*he*^ mice (Fig. 5a; Fig. S9), which was not observed in ISRIB-injected WT or *2b5*^*ho*^ animals.

**Fig. 5.**
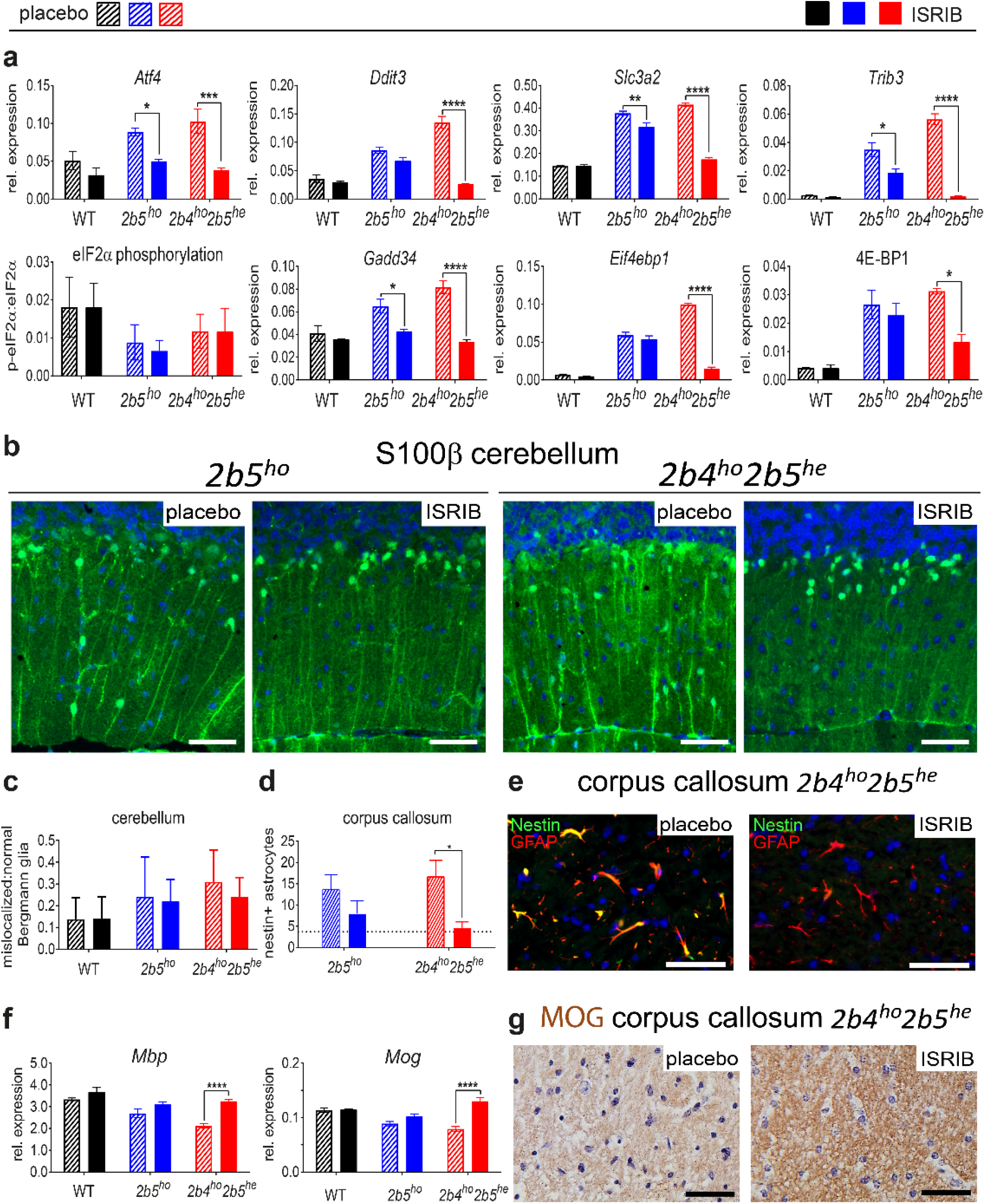
ISRIB modulates aberrant expression of ATF4-regulated mRNAs and ameliorates neuropathological astrocyte markers in VWM mouse cerebellum and corpus callosum. (**a**) Cerebellar expression of ATF4-regulated mRNAs, eIF2α phosphorylation and 4E-BP1 protein expression were measured in placebo- and ISRIB-treated WT and VWM mice (n=6 per group). Graphs show average +/-sd. Two-way ANOVA with Tukey’s correction was performed for each target. (**b**) Brain sections from placebo- and ISRIB-treated *2b5*^*ho*^ and *2b4*^*he*^*2b5*^*ho*^ mice (n=2 per group) were stained with antibodies against S100β. Thickness of Bergmann glia (BG) processes is reduced by ISRIB in *2b4*^*he*^*2b5*^*ho*^ mice; white bar, 0.05 mm. (**c**) Counts of mislocalized and normally localized S100β-positive Bergmann glia shows that ISRIB does not fully normalize Bergmann glia location in VWM mice. Differences in the ratio of mislocalized:normal localized Bergmann glia were statistically assessed by two-way ANOVA with Tukey’s correction. (**d,e**) ISRIB reduces number of nestin-GFAP double positive astrocytes in *2b4*^*he*^*2b5*^*ho*^ mice. The average number of nestin-GFAP double positive astrocytes in 4 untreated WT animals was included as reference (indicated as dotted line in the graph). Statistical differences were determined with one-way ANOVA using Tukey’s correction. White bar in immunofluorescence, 0.05 mm. (**f**) ISRIB restored levels of mature myelin mRNA markers in *2b4*^*he*^*2b5*^*ho*^ mice. Graphs show average +/-sd (*Akt*+*Gapdh*, qPCR reference). Two-way ANOVA was performed for each target using Tukey’s correction. (**g**) ISRIB restores immunoreactivity for mature myelin protein MOG in white matter structures in *2b4*^*he*^*2b5*^*ho*^ mice. Black bar, 0.05 mm. For all graphs, *p<0.05, **p<0.01, ***P<0.001, ****p<0.0001.

Bergmann glia localization in VWM mouse brain was improved by ISRIB, as visualized by S100β IHC, more clearly in *2b4*^*ho*^*2b5*^*he*^ than in *2b5*^*ho*^ mice (Fig. 5b). In quantification the changes in proportion of mislocalized Bergmann glia did not reach significance (Fig. 5c). Bergmann glia were localized closer to the Purkinje cells with ISRIB treatment, but did not reach or maintain their normal position completely (Fig. 5b).

The number of nestin and GFAP double-positive immature astrocytes normalized in ISRIB-injected *2b4*^*ho*^*2b5*^*he*^ mice and was reduced in *2b5*^*ho*^ mice, although statistical significance was not reached for the latter genotype (Fig. 5d, e). ISRIB effects on myelin in VWM mice were evident in mature myelin MBP and MOG mRNA levels and MOG IHC only in *2b4*^*ho*^*2b5*^*he*^ mice (Fig. 5f, g). No ISRIB effects on histology were observed in WT mice (data not shown).

The differential effect of ISRIB on body weight, clinical signs, expression of the ATF4-regulated transcriptome and histopathology in *2b5*^*ho*^ mice versus *2b4*^*ho*^*2b5*^*he*^ mice is striking (Fig. 4 and 5). ISRIB concentrations in venous blood samples drawn after 2 weeks of injections were similar in *2b5*^*ho*^ and *2b4*^*ho*^*2b5*^*he*^ mice, albeit lower than in WT mice (Fig. S10a). ISRIB’s concentrations in post-mortem blood and brain samples were similar in all three genotypes (Fig. S10b), indicating that ISRIB’s differential effect on phenotypic measures cannot be attributed to differences in ISRIB bioavailability in brain. ISRIB’s therapeutic effect may be influenced by the identity of the mutation. A recent study shows that ISRIB’s efficacy is increased by the Arg483Trp mutation in human eIF2Bδ (*19*). We measured ISRIB’s effect on protein synthesis rates in cultured astrocytes from WT, *2b5*^*ho*^ and *2b4*^*ho*^*2b5*^*he*^ mice undergoing a UPR (Fig. S10c). ISRIB was more effective in *2b4*^*ho*^*2b5*^*he*^ astrocytes than in WT and *2b5*^*ho*^ astrocytes, although statistical significance was not reached (Fig. S10c). In parallel, we determined ISRIB’s effect on protein synthesis in transfected astrocytes and found that protein synthesis in *2b4*^*ho*^*2b5*^*he*^ astrocytes was more sensitive to low concentrations of ISRIB than WT astrocytes (Fig. S10d). Increased ISRIB sensitivity was not observed in *2b5*^*ho*^ astrocytes (Fig. S10d).

## Discussion

VWM is a devastating leukodystrophy characterized by disappearance of cerebral white matter (*1*). In 2001, it was found that the basic VWM defect concerns eIF2B, which is pivotal in initiating translation and regulating its rate, especially under ISR activating conditions (*4, 12, 39, 40*). The housekeeping function of eIF2B is in stark contrast with its selective brain white matter involvement. Until now, the pathomechanisms of VWM have remained unclear. Available evidence indicates primary astrocyte involvement, secondary impediment of oligodendrocyte maturation and function, and negligible neuronal involvement (*3, 10, 41, 42*).

In this study we aim at unraveling VWM pathophysiology. Using polysomal profiling to reveal mRNAs that are differentially translated in brains of *2b5*^*ho*^ compared to mice, we demonstrate increased expression of an ATF4-regulated transcriptome in *2b5*^*ho*^ brain from an early age, before disease onset. Its expression level correlates with disease development and severity. Its expression in *2b5*^*ho*^ mice parallels the appearance of nestin+ immature astrocytes in the corpus callosum (Dooves 2016). Strikingly, this ATF4-regulated transcriptome is expressed selectively in astrocytes and not in other cells in VWM brain. We show that this astrocyte-selective expression is specific for VWM and is not a general response to neurodegeneration.

Previously, activation of the UPR was hypothesized in VWM, presumably through a feedback response sensitizing the ER to stress (*26*). Our current study disproves this concept. Differential expression of ATF6/XBP1s-regulated mRNAs was not detected in VWM patients. The reduced levels of phosphorylated eIF2α in brains of *2b5*^*ho*^ mice and VWM patients argues against cell stress and UPR activation as initiating cause for the enhanced ATF4 transcriptome in VWM astrocytes. Reduced activity of mutant eIF2B (*12*) by itself can explain the increased expression of ATF4 and its transcriptome. Increased GADD34 can explain the decreased levels of phosphorylated eIF2α. Reduced eIF2α phosphorylation probably compensates, albeit partially, the reduced activity of the mutant eIF2B. If the compensation had been sufficient, ATF4 levels would have been similar in WT and mutant brain tissue.

Results obtained with ISRIB indicate that the ATF4-regulated transcriptome is an essential part of VWM pathomechanisms. We previously demonstrated that guanabenz improves white matter and astrocyte pathology in *2b5*^*ho*^ mice (*43*). Guanabenz is an α2-adrenergic receptor agonist that also targets the ISR and reduces expression of the ATF4-regulated transcriptome (*44*). We now demonstrate that ISRIB, which activates eIF2B and leaves the α2-adrenergic receptor unaffected, reduces the expression of the ATF4-regulated transcriptome, improves VWM astrocyte and white matter pathology and ameliorates the clinical disease in VWM mice. Based on our findings, we conclude that reduced activity of mutant eIF2B initiates the disease in VWM. It activates the enhanced expression of ATF4 and ATF4-regulated transcriptome, which is moderated through the GADD34 feedback loop, decreasing eIF2α phosphorylation, albeit insufficiently to normalize eIF2B activity and prevent the disease.

Surprisingly, ISRIB did not affect eIF2 phosphorylation in VWM mice in our experiment. An increase in eIF2 phosphorylation would have been expected, as ISRIB reduced *Gadd34* mRNA levels in VWM mice. Possibly, the ISRIB-induced reduction in GADD34 protein level was not sufficient to achieve this. The ATF4 expression was not completely normalized by ISRIB in either VWM genotype, as exemplified by 4E-BP1 protein expression that was still detected in their brain tissue (Fig. 5). Both findings suggest that ISRIB treatment does not fully normalize the eIF2B activity, but leads to a better compensated system that still in part depends on reduced eIF2 phosphorylation.

ISRIB stabilizes the eIF2B complex formation through an interaction with the eIF2Bδ:eIF2Bβ interface (*20, 46*). Its effect may thus be influenced by the specific VWM mutation. The eIF2Bδ Arg483Trp mutation interferes with eIF2B complex stability (*12*), whereas the eIF2Bε Arg195His mutation does not and possibly reduces eIF2B activity through another mechanism (*48*). Our results indicate that ISRIB may preferentially impact eIF2B with complex-destabilizing mutations and warrant further investigation of the effect of specific mutations on eIF2B complex formation and activity.

The question how ISR deregulation causes pathology remains difficult to answer. Our overrepresentation analyses on the regulated mRNAs in brain of *2b5*^*ho*^ mice highlight increased functions of amino acid transport and biosynthesis of serine, glycine and cysteine. Several of these genes are regulated by ATF4 (*13, 44*). Increased serine and glycine levels have been reported in cerebrospinal fluid in VWM patients (*49*), indicating that these processes are indeed deregulated. Interestingly, serine, glycine and cysteine are used with glutamate for glutathione (GSH) synthesis (*50*). The ATF4-regulated transcription is known to aid in counteracting tissue damaging reactive oxygen species (ROS) (*51*). ROS is one of the stressors increasing the level of eIF2α phosphorylation (*52, 53*), but in brains of VWM mice and VWM patients, eIF2α phosphorylation is low, suggesting that ROS levels are not abnormally enhanced in VWM brain. Thus, the increased synthesis of the antioxidant GSH in the absence of an increased ROS in astrocytes may induce reductive shifts in the steady-state redox potential of the VWM brain (*50, 54, 55*). Especially in astrocytes a reductive shift may interfere with proliferation and maturation of oligodendrocyte precursor cells as reducing molecules have been shown to interfere with oligodendrocyte maturation (*56-58*). Strikingly, an astrocyte-dependent impairment of oligodendrocyte maturation is a central feature of VWM (*3, 10, 41, 59*). The impact of astrocytic redox imbalance may be higher on oligodendrocytes than on neurons, which intrinsically have extremely high levels of glutathione (*60*), explaining the sparing of this cell type. All data combined suggest that the defect in oligodendrocyte maturation may be due to a reductive shift in redox potential in mutant astrocytes effectuated by the expression of ATF4 (Fig. 6).

**Fig. 6.**
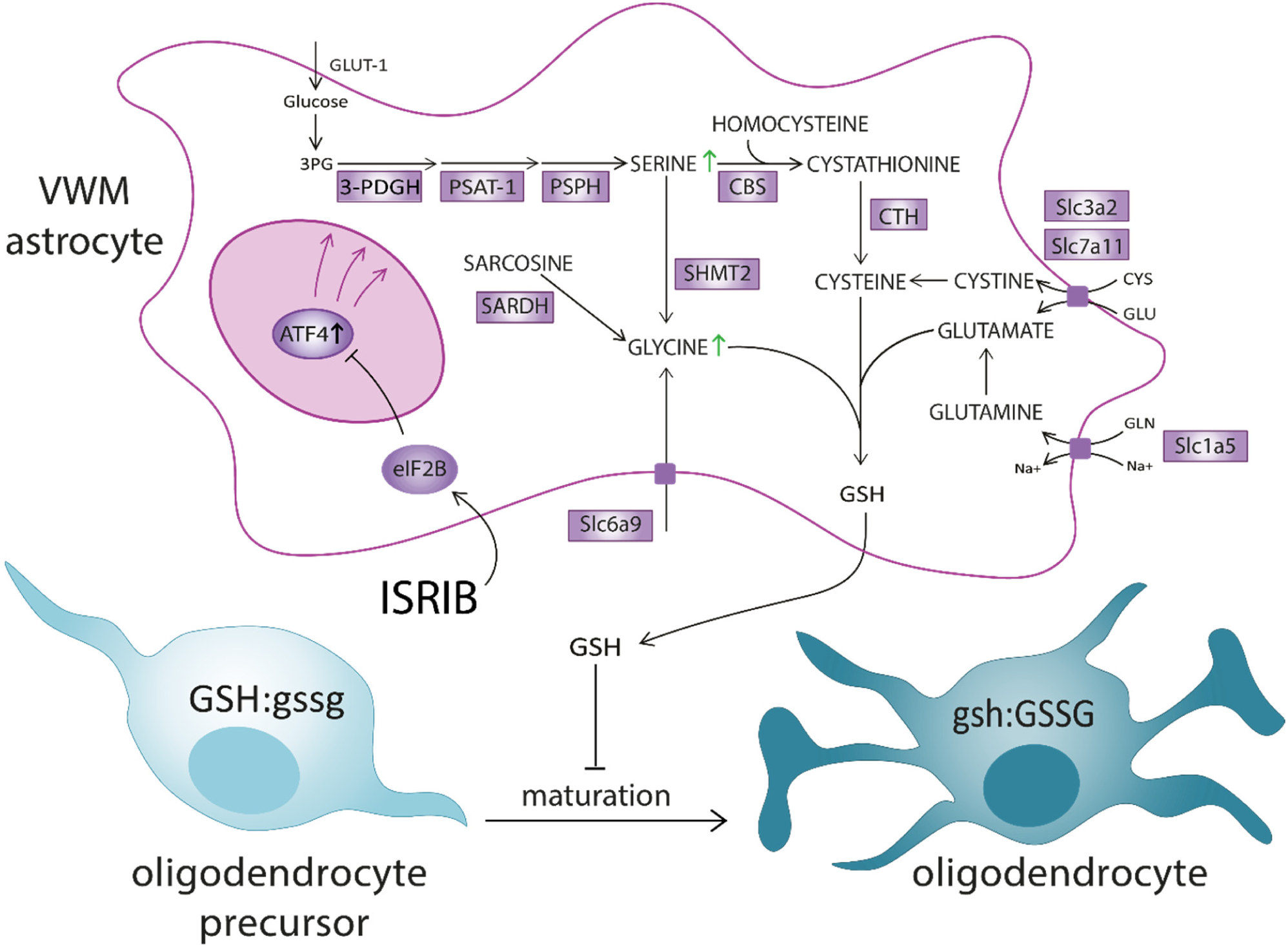
Molecular model for failing oligodendrocyte maturation in VWM and molecular effects of ISRIB. ISR-deregulation in VWM astrocytes increases ATF4, which regulates expression of mRNAs (indicated are upregulated mRNAs in magenta-shaded boxes). The ATF4-regulated transcriptome in VWM astrocytes affects amino acid metabolism, serine biosynthesis and glycine metabolic processes resulting in increased levels of the reducing compound glutathione (GSH). GSSG:GSH indicates redox potential (oxidized:reduced glutathione), with capitalized font indicating the most abundant type of glutathione (GSH, reduced; GSSG, oxidized). GSH may induce a reductive shift in redox potential interfering with maturation of oligodendrocyte precursors (OPC) to mature oligodendrocytes. The resulting reductive change in redox status could inhibit oligodendrocyte maturation, a process known to be driven by oxidative shifts in redox potential. Increased levels of serine and glycine (indicated with green arrows) have been described in cerebrospinal fluid from VWM patients. ISRIB impacts eIF2B activity and reduces ATF4 expression, consequently expecting to influence redox potential and improve OPC maturation.

There are several limitations to this study. We did not investigate if ISRIB suppresses VWM pathology and clinical signs for a full mouse life. These studies are long and licenses are difficult to obtain from ethical review boards, as they involve daily injections for several years. A more important limitation is that ISRIB efficacy has not been tested in humans. ISRIB targets eIF2B from various species, ranging from rabbit, hamster, mouse to humans (*16-20, 45-47*), suggesting that ISRIB is also efficacious in humans. An important limitation of ISRIB is that its effect is likely influenced by the specific VWM mutation, as indicated by our study, making that it is not equally effective in all VWM patients. Testing ISRIB on patient cells before ISRIB is prescribed may be required. An essential hurdle before studies can be undertaken to confirm efficacy in human patients is investigating the bioavailability of the compound in humans, which may involve cumbersome optimization of the vehicle. Alternatively, a more soluble ISRIB derivative or an alternative ISR-impacting compound can be envisioned that is capable of traversing the blood-brain-barrier (*61, 62*).

### Concluding remarks

The role of ISR regulation in health and disease is currently at the center of attention in aging and various neurodegenerative diseases, including Alzheimer disease, Parkinson disease and ALS (*63-65*). This study is first in showing that modulation of ISR deregulation ameliorates neurological dysfunction in VWM mice without affecting mouse general health. We show that an ISR deregulation selectively occurs in brain astrocytes of VWM mice and VWM patients. Effects of ISR modulation demonstrate that its derangement is a pivotal part of VWM pathogenesis. ISRIB reduces the expression of ATF4-regulated transcriptome, improving disease outcome in VWM mouse models. The study reveals that the ISR is a viable drug target for treating VWM.

## Materials and Methods

### Animal models

The mouse strains with a homozygous point mutation in the eIF2Bε (*Eif2b5*^*Arg191His/Arg191His*^, *2b5*^*ho*^) or eIF2Bδ subunit (*Eif2b4*^*Arg484Trp/Arg484Trp*^, *2b4*^*ho*^) are described elsewhere (*10*). Crossbreeding of the two mutant lines generated mice that are homozygous for the eIF2Bδ Arg484Trp mutation and heterozygous for the eIF2Bε R191H mutation (*2b4*^*ho*^*2b5*^*he*^) or heterozygous for the eIF2Bδ Arg484Trp mutation and homozygous for the eIF2Bε R191H mutation (*2b4*^*he*^*2b5*^*ho*^). Mice were weaned at P21 and kept with a 12 hour light/dark cycle with food and water provided *ad libitum*. The animals were sacrificed at different ages depending on the experiment. When the mutant mice became so ataxic that they could not reach the food on top of the cages, food was placed inside the cages and eventually replaced by suspensions of water and ground food pellets. The humane end point (HEP) for the mutant mice was defined by weight loss of more than 15% of body weight for 2 consecutive days and severe ataxia.

### Study approval

All animal experiments were carried out in compliance with the Dutch and European law and with approval of the local animal care and use committee of the VU University [license FGA 11-05A5]. Written informed consent was provided by patients or their parents for the collection and use of patients’ samples. The Medical Ethics Committee of the VU University Medical Center approved of the procedures.

### Sample preparation

Mice were sacrificed by cervical dislocation at indicated ages. The individual organs were snap-frozen in liquid nitrogen immediately after they were taken out. Brains were divided in four parts: 1) olfactory bulb (which was discarded and not investigated), 2) forebrain including cortex and corpus callosum, 3) cerebellum and 4) the remaining structures, including striatum and hippocampi. All organs were stored in a −80°C freezer until further use, except for obtaining nuclear lysates which were prepared from fresh tissue.

Cytoplasmic lysates of mouse organs were prepared by grinding each organ with pestle and mortar under liquid nitrogen. All chemicals were purchased from Sigma-Aldrich unless otherwise stated. The frozen powder was resuspended with a minimixer in 600 μl lysis buffer, composed of 20 mM Tris pH 7.4, 100 mM potassium acetate, 3 mM magnesium acetate, 3 mM dithiothreitol, 0.5% Triton-X100, 0.5% sodium deoxycholate and protease inhibitors (Roche). Samples were kept on ice for 15 min and triturated three times with a 23G needle. The samples were centrifuged for 10 min at 4 °C to remove nuclei, membranes and debris. Samples were aliquoted and stored at −80°C or applied to sucrose gradients for fractionation into polysomes and monosomes.

### Polysomal profiling

Equal volumes of cytoplasmic lysate were loaded onto a 15%-50% isokinetic sucrose gradient (containing 1 ug/ml cycloheximide, 100 ug/ml heparin, and 7 mM β-mercaptoethanol) and centrifuged for 45 min at 4°C in a Beckman SW55 ultracentrifuge at 52.000 rpm. The gradient was fractionated into polysome fractions and a monosome fraction. The gradients were pushed upwards by a 70% glycerol solution through an Uvicord with a velocity of 1 ml per min. mRNA was isolated from the fractions using Trizol™ reagent (Invitrogen), after addition of 300 mM sodium acetate and 20 mM EDTA. After mixing by inverting the tube, the solutions were frozen for at least 20 min at −80°C. After thawing chloroform was added and the sample was centrifuged for 10 min at 4000 rpm (4 °C; Eppendorf). The aqueous phase was isopropanol-precipitated on ice in the presence of linear acrylamide (20 µg/ml; Invitrogen). The pellet was air-dried for approximately 1 min and resuspended in non-DEPC-treated nuclease-free water (Ambion). Total RNA was precipitated with LiCl (4 M final concentration) at −20°C for at least one hour. Samples were centrifuged at maximum speed (Eppendorf) for at least one hour at 4 °C. The pellet was washed with ice-cold 70% EtOH, resuspended in 20 μl non-DEPC-treated nuclease-free water (Ambion). An additional ethanol precipitation was performed if the A230/A260 ratio of the RNA was suboptimal (measured by (NanoDrop 2000C, ThermoFisher). mRNA was subjected to Cy3 and Cy5 labeling and labeled cRNA was applied to expression microarray analyses at the local microarray facility using Agilent 4x44K *Mus musculus* dual channel array slides, each consisting of 4 arrays with 44,000 2-channel probe spots (Cy3 and Cy5).

### Data processing and variance analysis

Expression data were obtained from two independently conducted microarray experiments, each using two mice per genotype. In the first experiment the gradient was fractionated into one polysome (B) and one monosome fraction (A) and in the second experiment into two polysome fractions (B1, B2) and one monosome fraction (A). The signals for mRNAs in B1 and B2 fractions were averaged. Parsing and normalization of the microarray data was done in R using the Limma package (*66*). Normalization was done in two steps, firstly within-array normalization was applied in the form of Loess normalization without background subtraction. Secondly, between-array normalization was applied using quantile normalization. Variance was evinced through estimation of within-block correlation using the intraspotCorrelation function and subsequently by fitting linear models to all probe log-intensities, using the lmscFit function (*67*). Two different kinds of variance were estimated, firstly the variance in mRNA abundance between WT and *2b5*^*ho*^ was assessed and secondly the variance in the polysome densities of each mRNA between WT and *2b5*^*ho*^ was calculated. The former was done by taking the difference of ^2^log-transformed expression values: Δ(Δ(B_*2b5ho*_,A_*2b5ho*_),Δ(B_wt_,A_wt_)) and Δ(B_*2b5ho*_,B_wt_), in which B refers to polysome fraction and A to monosome fraction. These tests are referred to as ΔΔBA and ΔB respectively. For all tests a false discovery rate (FDR) cut-off of 0.05 was calculated according to the Benjamini-Hochberg method (*68*) to select probes that show reliable variance.

### Overrepresentation analysis

A threshold was set for what constitutes gene expression in the resulting dataset. The probe average intensity was plotted against its frequency, showing a bimodal distribution of expressed and non-expressed genes. The probe intensities were plotted against the false discovery rate (FDR) to determine the threshold between expression and non-expression. The intensity for which FDR was smaller than 0.05 was 6.29; this intensity was taken as the cut-off value for what constitutes expression. Using this cut-off a denominator list was generated, representing WT mouse brain tissue expression. This list consists of 22,473 probe sets, out of a total of 43,379. Next, we generated a list of probe sets that are differentially associated with polysomes from WT versus *2b5*^*ho*^ brain (ΔB). Overrepresentation analysis was performed with the software program Go-Elite version 1.2.5 (*69*). In short, the list of regulated mRNAs (ΔB) in *2b5*^*ho*^ mouse brain was investigated for pathway, ontology or gene set. The number of permutations was kept at the default value of 2000. Only results being non-redundant with a Z-score> 2, a p-value< 0.05, a FDR<0.2 and fold change of > 1 or <-1 were selected.

To investigate enrichment of ATF4-regulated mRNAs we compared published data sets of ATF4-dependent mRNAs (Suppl. Table S4 in (*13*)) with the denominator list as well as the differentially polysome-associated mRNAs (ΔB). Significant differences were determined using a Fisher exact-square test. A similar approach was undertaken for a microarray analyses on total RNA samples from postmortem human white matter tissues (6 controls, 6 VWM patients, details listed in Supplementary Data File 4). A table of homologous genes of mouse and human was downloaded on January 16, 2018 from the <u>Mouse Genome Database (MGD)(</u>*<u>70</u>*). The “Symbol” column in this table was used to match human genes from the microarrays of this study with the mouse genes from the published study (*13*). Enrichment was subsequently determined by comparing the same published data set of ATF4-dependent mRNAs with the denominator list (cut-off determined as 6) and the differentially accumulated mRNAs (FDR<0.05). In parallel from the same publication we distilled ATF6- and IRE1α-regulated mRNAs by subtracting ATF4-regulated mRNAs from the UPR-regulated mRNAs in response to tunicamycin (Suppl. Table S4 (*13*)). This non-ATF4 list was then compared with the denominator list (cut-off determined as 6) and the differentially accumulated mRNAs (FDR<0.05) in VWM white matter. Significant differences were determined using a Fisher exact-square test.

### Quantitative PCR (qPCR)

Total RNA was extracted from mouse brain and human white matter with Trizol™ reagent (Invitrogen). Reverse transcription and qPCR were performed as described previously (*71*). Primers were designed using PerlPrimer v1.1.1928 or PrimerBLAST. Depending on the experiment *Akt, Gapdh* or *Akt* + *Gapdh* mRNA was used as reference as indicated in figure legends. *Gapdh* mRNA is not a suitable reference for the qPCRs on polysomal profiling: *Gapdh* mRNA is so efficiently translated that it is found abundantly in B2 fraction and much less so in A and B1 fractions ((Fig. S2). *Akt* mRNA has a consistent abundance in all gradient fractions in all tested genotypes (data not shown). In contrast, *Akt* mRNA abundance in WT and *2b5*^*ho*^ mouse brain does not differ, but declines from postnatal day 0 to 14 after which it remains stable (data not shown). *Gapdh* mRNA abundance in brain is consistent during mouse life and does not differ between WT and VWM genotypes. The primers used in this study are listed in Suppl. File 1.

### Western blotting

Total protein concentration in cytoplasmic was measured by Bradford according to the manufacturer’s instructions. Equal amounts of protein were applied to SDS-PAGE. In-gel protein loading and sample transfer was checked with 2,2,2-trichloroethanol (TCE) as described (*72*). Membranes were blocked with 5% non-fat-dry-milk and incubated with one of the following primary antibodies (anti-eIF2α, SC-11386/SantaCruz or anti-4E-BP1, 53H11/Cell Signaling, or anti-ATF4 D4B8/Cell Signaling). Membranes used for detecting phosphorylated eIF2α were blocked with 5% bovine serum and incubated with anti-Ser51-phosphorylated eIF2α antibody (Cell Signaling, D9G8). After incubating with secondary antibody (HRP-labeled anti-IgG rabbit, 1:10000, Dako, P0448), the signal was detected with an enhanced chemiluminescent substrate (SuperSignal West Femto Substrate, Fisher Scientific) and imaged with Image Studio on the LI-COR Odyssey Fc Imager (*71*).

### Immunohistochemistry

Immunohistochemistry (IHC) and immunofluorescence (IF) was performed as described(*10*). IHC and IF were performed with antibodies against ATF4 (#11815, 1:50, Cell Signaling), 4EBP1 (#9644, 1:500, Cell Signaling), GFAP (Z0334, 1:1000, DAKO), Olig2 (AB9610, 1:200, Millipore), S100β (S2532, 1:1000, Sigma), nestin (#611658, 1:200, BD Bioscience). Briefly, brains were perfusion-fixed with 4% paraformaldehyde (PFA) and embedded in paraffin. Paraffin-embedded tissue sections (6-μm-thick) were deparaffinized and endogenous peroxidases where blocked with 0.3% H_2_O_2_. After heat-induced antigen retrieval in 0.01 M citrate buffer (pH 6) or 10 mM Tris, 1 mM EDTA buffer (pH 9), sections were blocked with 5% normal goat serum (Thermofisher Scientific) for S100β this blocking step was replaced by a Mouse on Mouse (M.O.M.™) blocking reagent (Vector laboratories). For ATF4 detection, tissue was permeabilized with 0.1% saponin and 0.5% Triton X100. Immunoreactivity was detected with DAB chromogen (DAB+, DAKO). For double stainings, tissue was incubated with primary antibody and developed with 3,3′-diaminobenzidine DAB as described above. After a second heat-induced antigen retrieval step, incubation of the second primary ab was performed. Immunoreactivity was detected with liquid permanent red (LPR) substrate chromogen (DAKO). Images were taken with a Leica DM6000B microscope (Leica Microsystems). Numbers of nestin-positive astrocytes were determined by immunofluorescence as described (*10*). The ratio of mislocalized:normal Bergmann glia was determined basically as described(*10*) with minor changes: secondary antibodies against anti-IgG were labelled with 488 (S100β) rather than HRP and counterstained with 0.1% Sudan Black. Images were obtained with a Leica 5000b fluorescent microscope. DAPI was included in mounting media to stain for nuclei.

### ISRIB trial

A vehicle for solubilizing ISRIB was developed to ensure ISRIB delivery from the intraperitoneal cavity to the blood and tissues (SEPS Amatsi). A 1 mg/ml stock solution was prepared with polysorbate 20 as surfactant and diluted 10-fold with water for injection (WFI) immediately prior to injection. The same vehicle alone without ISRIB was made in parallel as placebo and diluted 10-fold with WFI as well. 1 mg/kg ISRIB was injected daily into male WT, *2b5*^*ho*^ mice and *2b4*^*ho*^*2b5*^*he*^ mice. In parallel, mice with either of three genotypes (WT, *2b5*^*ho*^ and *2b4*^*ho*^*2b5*^*he*^) were injected with the solubilizing vehicle only (placebo) to investigate side-effects caused by the daily injection procedure and constituents of the vehicle. Group sizes were n=6 for WT mice and n=14 for VWM mice (n=68 in total). Injections were placed on either left hand or right hand side of the abdominal wall at alternating days. Body weight was measured daily and blood samples were taken by submandibular vein bleed after two weeks of daily injections.

ISRIB concentrations were determined in venous blood two weeks after start of the injections and in post-mortem blood and brain as previously described (*15*) with some minor modifications. ^2^H_10_-ISRIB (synthesized by Herman ten Brink, Amsterdam UMC, location VUmc) was added to the samples for precise quantification. The samples were reconstituted in acetonitrile and separation of ISRIB was achieved on a Waters Acquity UPLC BEHC18 1.7µm 2.1x100mm column and quantitative analysis was performed by LC-MS/MS by using an API 5000 (AB Sciex) mass spectrometer in positive electrospray ionization mode with MRM m/z 451-266 and m/z 461-276 for ISRIB and ^2^H_10_-ISRIB respectively.

Motor skills (neuroscores, Balance Beam) were assessed at indicated time points as previously described (*10, 73*). Mice were subjected to a catwalk set-up for a quantitative gait analysis, using CatWalk XT 10.6 (Noldus Information Technology) prior to being euthanized. The machine consists of an enclosed glass walkaway (width), illuminated from the distal long edge with a green light and from the ceiling with a red light. Mice are placed onto the glass plate and as the animals’ paws get in contact with the glass floor, the light is internally reflected, captured by a high-speed video camera and transformed in a digital image. Images were analysed by using the automated footprint classification provided in the CatWalk XT 10.6 software with a minimal of consecutive steps of six, average speed ranging from 20.0 to 60.0 cm/second and a maximum allowed speed variation of 40%. All CatWalk data have been analysed by the same researcher in order to reduce interrater variabilities and further processed by a different researcher, blinded to treatment and genotype (*74*). At the end of the experiment animals were euthanized by cervical dislocation or perfusion-fixation with 4% PFA. Brains were taken out for molecular and neuropathological examination.

### Protein synthesis measurements

Protein synthesis rates were measured in adult mouse astrocytes with AHA-incorporation and statistical analysis of the data was performed as previously described (*75*). Primary cultures of adult mouse astrocytes were transfected with pGL3-*Gapdh and* pNL1.1-*Atf4* as described (*75*). Cells were exposed to 300 nM thapsigargin for 16 hours in absence or presence of increasing concentrations of ISRIB. Luciferase activity was measured as before. The pNL1.1-*Atf4* construct comprises the murine *Atf4* promoter from nucleotide −1600 to +1023 with respect to the transcription start site (the intron 1 sequence is not present in the construct). The construct was made as described (*75*) using infusionforward primer (5’-TGGCCTAACTGGCCGCACTGGCTGTTTGGAGAC) and infusionreverse primer (3’-AGTGTGAAGACCATGTTGTGGGGCTTTGCTGGA). The construct was not responsive to thapsigargin treatment nor ISRIB and therefore the pNL1.1-*Atf4-*derived luciferase activity was not included in the manuscript.

### Statistical analysis

For the validation of polysome association using qPCR, statistical differences were determined using a two-way ANOVA (genotype and fraction type as factors) followed by a Sidak’s multiple comparison test, according to GraphPad Prism software.

Comparisons between WT and *2b5*^*ho*^ samples were analyzed with Student’s t-test using GraphPad Prism software. For qPCR and Western blot analysis between more than two genotypes, statistical differences were determined using a one-way ANOVA followed by a Tukey’s multiple comparison test, using GraphPad Prism software. Differences were interpreted as significant when p<0.05.

Statistical analysis on the differences at groups of different ages was done for each mRNA using a two-way ANOVA (genotype*day interaction all days) followed by 2 two-way ANOVAs for each mRNA (one comparing data of <P21 with ≥P21 and one comparing data of <P28 with ≥P28 for genotype*day interaction) using SPSS; p<0.01 was considered significant.

Statistical analysis investigating the ISRIB differences within WT, *2b5*^*ho*^ and *2b4*^*ho*^*2b5*^*he*^ genotypes in body weight was performed with a repeated measurements two-way ANOVA. Statistical analyses investigating the ISRIB differences within and among WT, *2b5*^*ho*^ and *2b4*^*ho*^*2b5*^*he*^ genotypes in ISRIB concentrations, Balance Beam, CatWalk parameters, qPCR, Western blot, ratio of mislocalized to normally localized (normal) S100β Bergmann glia were performed with a two-way ANOVA with a post Tukey’s multiple comparison test. Statistical analysis investigating ISRIB differences in numbers of nestin-positive astrocyte within *2b5*^*ho*^ and *2b4*^*ho*^*2b5*^*he*^ genotypes were statistically assessed with one-way ANOVA with a post Tukey’s multiple comparison test (untreated WT animals were included as reference).

Statistical differences between ISRIB sensitivity in WT2 vs *2b5*^*ho*^ and WT1 vs *2b4*^*ho*^*2b5*^*he*^ and WT1 vs WT2 adult mouse astrocytes were determined through a one-way ANOVA followed by Sidak’s multiple comparison test, using GraphPad Prism software. For the transfection assays, areas under curve (AUC) were calculated for each individual experiment after correcting for differences between experiments but not between test conditions within experiments (genotype, treatments) (*76*).

Results of all statistical analyses in the presented work are listed per figure in Supplementary Data File 5; for CatWalk data these results are listed in a separate tab. Differences were interpreted as significant when p<0.05, unless indicated differently.

## Acknowledgements

We thank Gerbren Jacobs for mouse genotyping. We thank Dave Speijer for fruitful discussions and critically reading the manuscript. We thank Nienke Wevers, Carola G. M. van Berkel, Gino Hu-A-Ng, Erwin E. W. Janssen, Herman ten Brink, Ilja Boor and Bastijn Koopmans for technical assistance. We thank Pim van Nierop for advice on overrepresentation analyses. We thank the staff of the VU mouse housing facility for mouse breeding and injections. We thank Maarten Loos for providing access to the CatWalk software. We thank Rogier Min for fruitful discussions during the preparation of the manuscript.

## Authors contribution

MSvdK conceptualized the polysomal profiling. AAMT and TEMA developed the protocol for sample and polysome preparation. TEMA and EP performed the polysomal profiling experiments. MB designed the microarray on human brain RNA. MJT and DM performed data analyses and variance analyses of microarray data of mouse brain and human brain. HF and LW performed overrepresentation analyses. LW, EP, NLP and TJtB performed the qPCR and Western blot experiments. LW, EJ performed the statistical analyses of data acquired with qPCR, Western blot. SvdS designed and advised on statistical methods. NLP performed IHC and IF experiments under supervision of MB. MSvdK supervised the use of human and mouse tissue. MV provided the *Gfap-Scap*^*-/-*^mice and advised on the experiment design. TEMA designed the ISRIB trial. TJtB, SB, NS, EJ and TEMA acquired and analysed data during ISRIB trial, including body weight, neuroscores, balance beam. NS performed CatWalk experiments and analysed the data. EAS designed and supervised the ISRIB measurements in blood and brain. LW and DVG performed the protein synthesis measurements and the statistical analyses of the acquired data. TEMA and MSvdK wrote the manuscript. TEMA and EJ made the figures. MSvdK and TEMA obtained funding. **Competing interest:** The authors have declared that no competing interest exists.

## Data and materials availability

Reagents (e.g. plasmids) and mouse models must be obtained through an MTA.

## Supplementary Materials

**Table S1.** eIF2Bε^Arg191His^–regulated shifts in mouse forebrain polysome association.

|  | Probe on<br>microarray chip | Gene | Transcript | FDR | logFC | Regulated by ATF4 <sup>1</sup> |
| --- | --- | --- | --- | --- | --- | --- |
| 1. | A_52_P532982 | *Gdf15* | NM_011819 | 6.13E-08 | 4.42 | - |
| 2. | A_51_P331570 | *Trib3* | NM_175093 | 4.27E-04 | 3.61 | + |
| 3. | A_51_P519251 | *Nupr1* | NM_019738 | 7.43E-04 | 2.87 | + |
| 4. | A_51_P453475 | *Slc1a5* | NM_009201 | 2.84E-03 | 2.62 | + |
| 5. | A_51_P269216 | *Atf5* | NM_030693 | 1.26E-03 | 2.53 | + |
| 6. | A_52_P374960 | *Ostn* | NM_198112 | 6.99E-03 | 2.40 | + |
| 7. | A_51_P263965 | *Hmox1* | NM_010442 | 3.84E-05 | 2.36 | + |
| 8. | A_51_P129502 | *Slc7a11* | NM_011990 | 3.91E-02 | 2.22 | + |
| 9. | A_51_P283473 | *Fibin* | NM_026271 | 2.67E-03 | 2.12 | + |
| 10. | A_51_P493987 | *Moxd1* | NM_021509 | 3.99E-04 | 2.03 | + |
| 11. | A_51_P421140 | *Tubb6* | NM_026473 | 1.73E-04 | 1.98 | - |
| 12. | A_51_P273639 | *Slc7a5* | NM_011404 | 9.75E-06 | 1.84 | + |
| 13. | A_52_P245648 | *Slc3a2* | NM_008577 | 3.91E-02 | 1.56 | + |
| 14. | A_51_P275016 | *Slc7a3* | NM_007515 | 2.64E-02 | 1.56 | + |
| 15. | A_51_P268697 | *Slc1a3* | NM_148938 | 2.30E-02 | 1.55 | - |
| 16. | A_52_P533146 | *Ddit3* | NM_007837 | 2.61E-03 | 1.51 | + |
| 17. | A_52_P21659 | *Pck2* | NM_028994 | 2.64E-02 | 1.45 | + |
| 18. | A_51_P319460 | *Osmr* | NM_011019 | 3.75E-02 | 1.32 | +<sup>2</sup> |
| 19. | A_51_P489153 | *Crot* | NM_023733 | 1.77E-02 | 1.28 | - |
| 20. | A_51_P427674 | *Cpt1a* | NM_013495 | 2.64E-02 | 1.19 | - |
| 21. | A_51_P317427 | *Psat1* | NM_177420 | 1.77E-02 | 1.17 | + |
| 22. | A_51_P427674 | *Cpt1a* | NM_013495 | 3.41E-02 | 1.05 | - |
| 23. | A_51_P136303 | *Cyp4f15* | NM_134127 | 2.64E-02 | 0.88 | - |
| 24. | A_52_P57013 | *Nxn* | NM_008750 | 4.24E-02 | -0.85 | - |
| 25. | A_51_P436269 | *Grin2c* | NM_010350 | 8.92E-03 | -1.14 | - |
| 26. | A_51_P209327 | *Apln* | NM_013912 | 2.64E-02 | -1.18 | + |
| 27. | A_52_P266686 | *Ntsr2* | NM_008747 | 4.56E-02 | -1.29 | - |
| 28. | A_51_P362066 | *Chi3l1* | NM_007695 | 2.85E-02 | -1.43 | - |
| 29. | A_52_P467930 | *BC158068* | BC158068 | 2.81E-02 | -1.47 | - |
| 30. | A_51_P194099 | *Thrsp* | NM_009381 | 2.64E-02 | -1.47 | - |
| 31. | A_51_P456465 | *Cldn10* | NM_021386 | 3.93E-03 | -1.57 | - |
| 32. | A_51_P244540 | *Ntsr2* | NM_008747 | 4.19E-04 | -1.72 | - |
| 33. | A_51_P443819 | *2610034M16Rik* | NM_027001 | 2.98E-02 | -1.91 | - |

<sup>1</sup>) reported in (13)
<sup>2</sup>) downregulation by ATF4 reported (13)

**Fig. S1.**
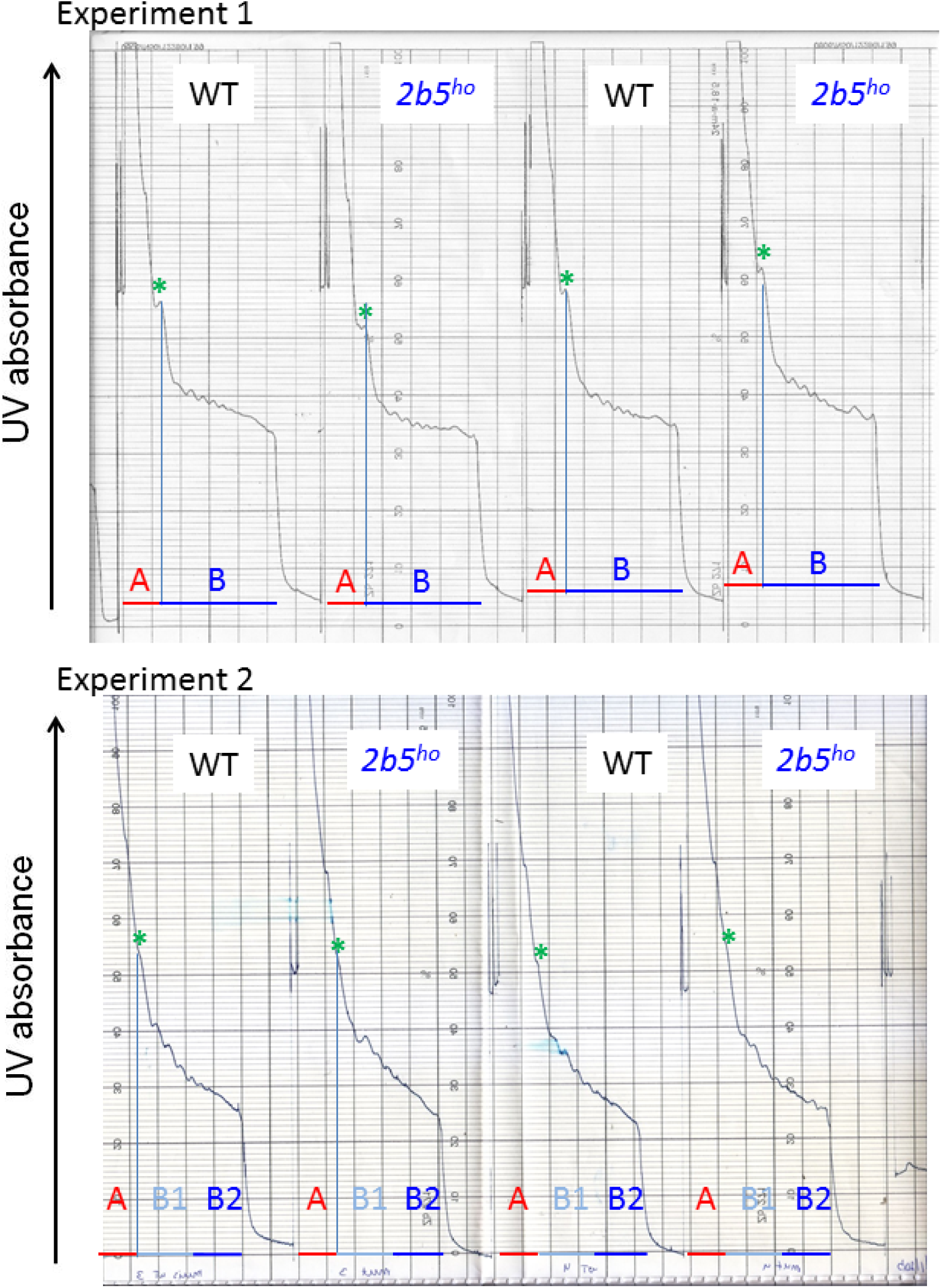
Polysome profiles of wild type and *2b5*^*ho*^ mouse brain are similar. Sucrose gradients were pushed upwards by a 70% glycerol solution through an Uvicord with a velocity of 1 ml per min. UV absorbance was measured and was the basis for the fractionation. Mouse genotypes are indicated. **\***, 80S ribosome; A, monosome fraction; B, polysomes; B1, polysomes with less than five ribosomes per mRNA; B2, polysomes with 5 or more ribosomes per mRNA.

**Fig. S2.**
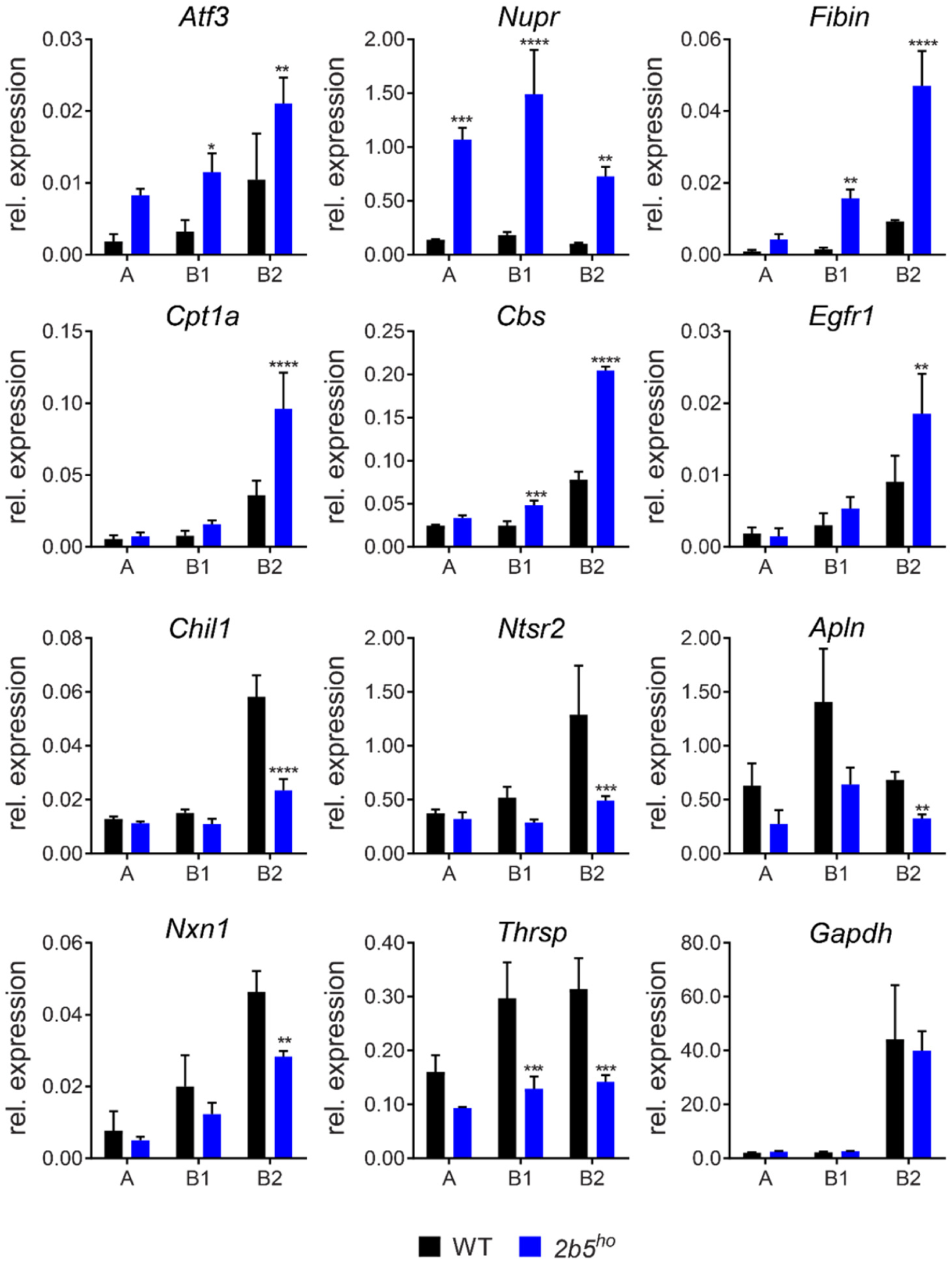
Verification of differentially associated mRNAs in *2b5*^*ho*^ mouse brain polysomes support ISR deregulation. The validated candidates are increased or decreased in polysome fractions from *2b5*^*ho*^ forebrain tissue compared to WT. mRNA levels were measured in the same samples as used in Fig. 1b of the manuscript (*Akt*, qPCR reference). Graphs show average ± sd (n=3). A, subpolysomal fraction; B1, polysomes with less than five ribosomes per mRNA; B2, polysomes with 5 or more ribosomes per mRNA. Statistical significance was determined by two-way ANOVA with Sidak’s correction, **p<0.01, ****p<0.0001.

**Fig. S3.**
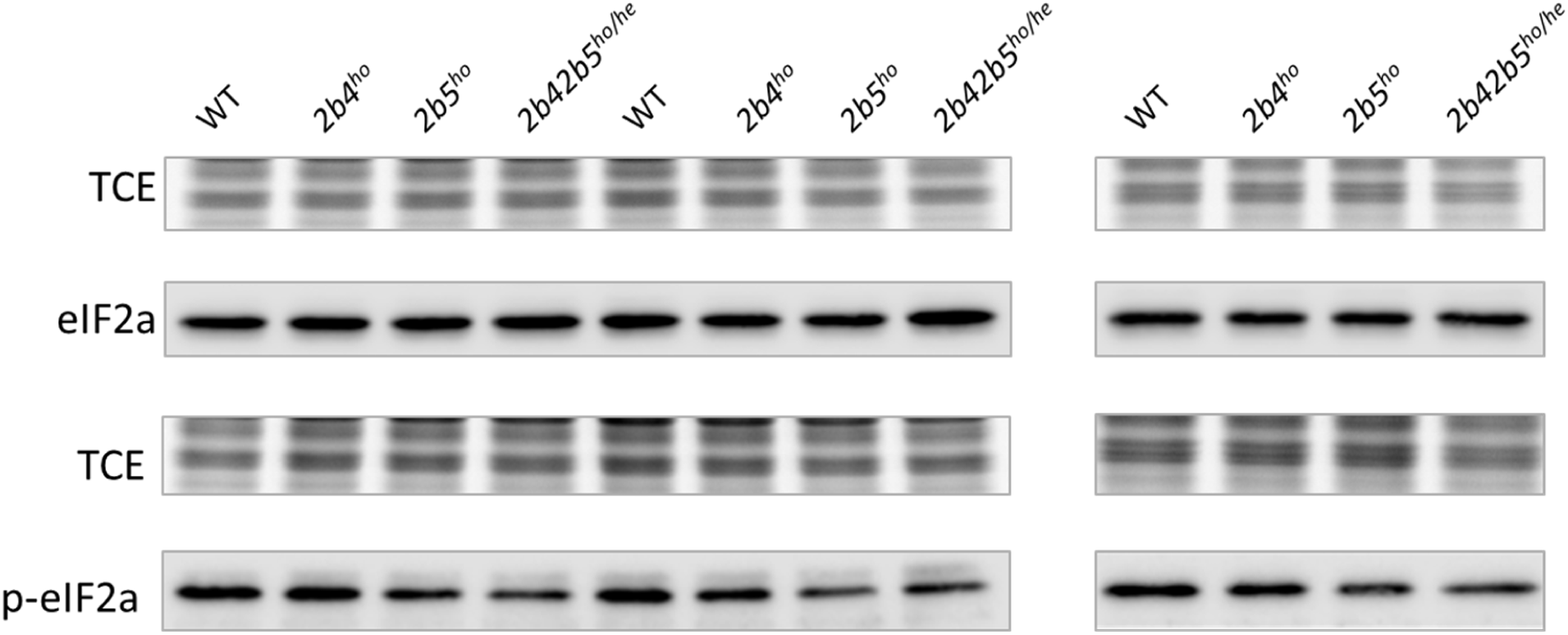
eIF2α phosphorylation is reduced in VWM mouse cerebellum. Equal amounts of protein were applied to SDS-PAGE. In-gel protein loading and sample transfer was checked with 2,2,2-trichloroethanol TCE) as described (*72*). Quantitative results are shown in Fig. 1c.

**Fig. S4.**
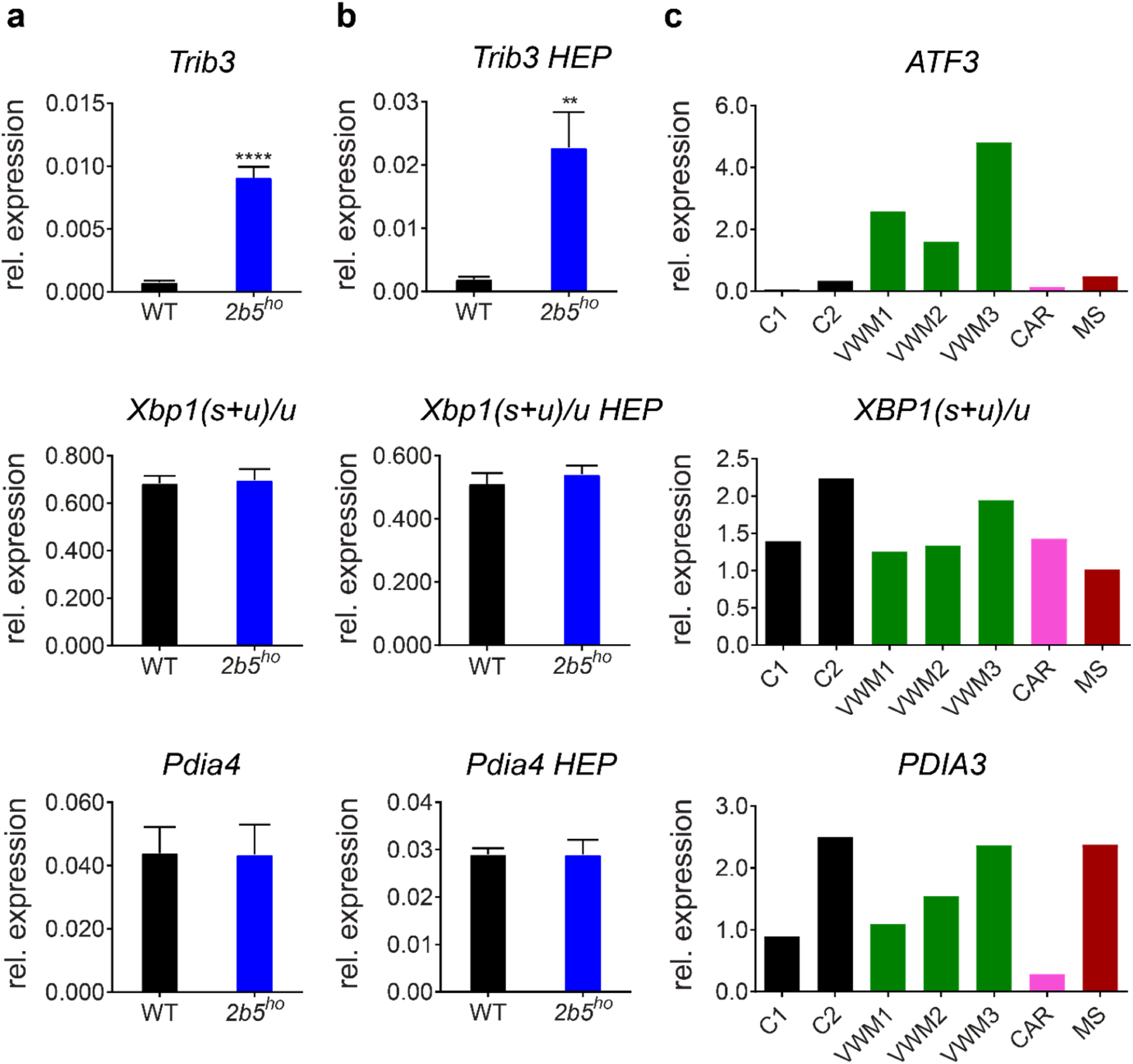
The ATF4-transcriptome is expressed in VWM mouse and VWM patients’ brain without markers indicating cellular stress. (**a**) mRNAs regulated by transcription factors ATF6 or IRE1α are similarly expressed in brains of 4-month-old WT and *2b5*^*ho*^ mice. *Pdia4* mRNA is regulated by ATF6 and *Xbp1* splicing (represented as *Xbp1(s+u)/u* ratio; s, spliced; u, unspliced) is regulated by IRE1α. *Trib3* was included as positive control for ATF4 regulation. (**b**) Similar expression levels were measured in brain of *2b5*^*ho*^ mice at humane end point (HEP). Graphs show average ± sd, n=6 for mRNA expression at 4 months (**a**) and n=3 at HEP (**b**). Comparisons between WT and *2b5*^*ho*^ samples are analyzed with Student’s T-test. **p<0.01, ****p<0.0001. (**c**) mRNA levels were quantified in postmortem frontal white matter tissue from VWM patients and controls (negative controls without brain pathology, C1, C2 and disease controls CAR and MS; CAR, CARASAL; MS, multiple sclerosis, *GAPDH*+*AKT* was used as qPCR reference). *ATF3* expression is regulated by ATF4 (ISR marker). *PDIA3* expression by ATF6 and *XBP1s* (represented as *XBP1(s+u)/u* ratio) by IRE1α (ATF6 and IRE1α are UPR-specific markers). *PDIA4* was not detected in human brain tissue. Each bar represents one measurement (n=1). Details on control and patients’ tissues are listed in Supplementary data file 4.

**Fig. S5.**
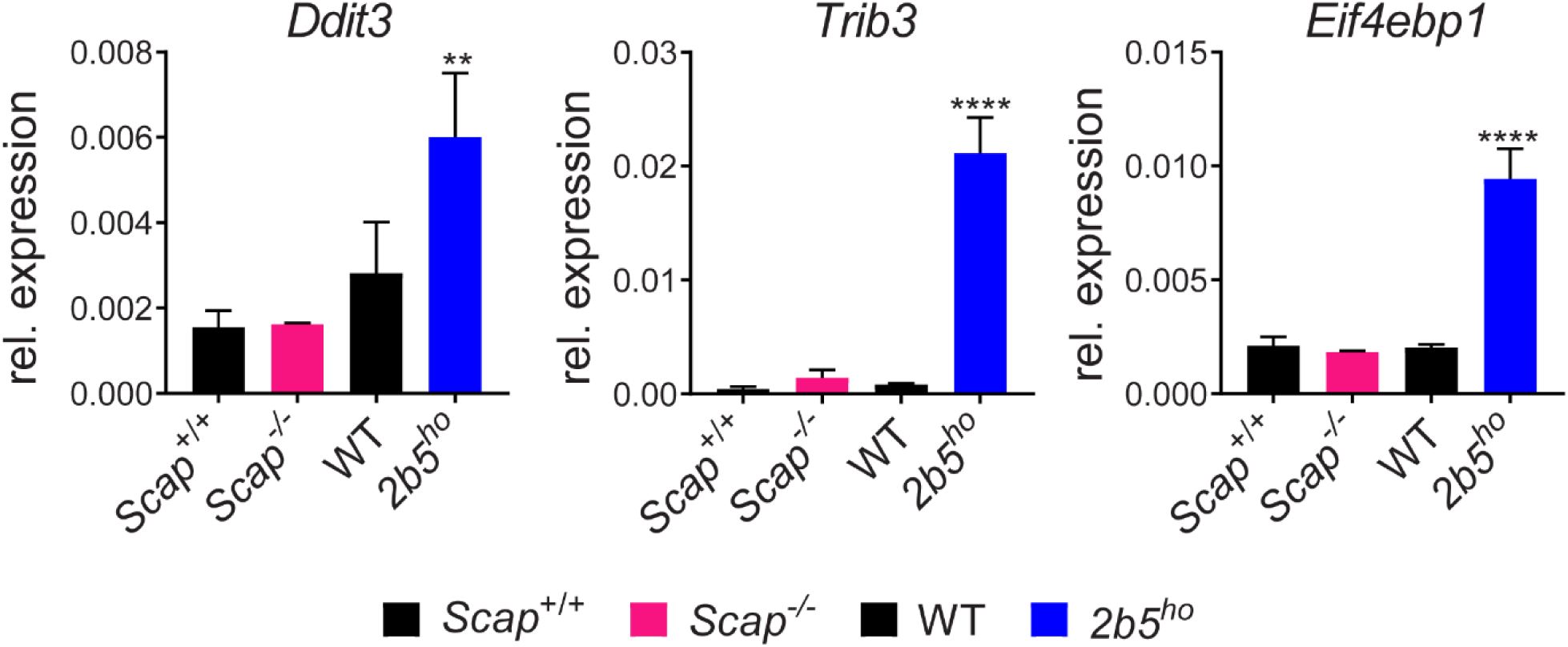
Enhanced expression of ATF4-regulated mRNAs is absent in brain tissue from *Gfap-Scap*^*-/-*^mice. mRNA levels were determined with qPCR (*Gapdh*+*Akt* was used as qPCR reference). *Ddit3, Trib3* and *EIf4ebp1* mRNA expression was measured in forebrain tissue of 2-month-old *Gfap-Scap-/-*mice (mid-stage disease) and compared to *2b5*^*ho*^ mice (each mutant mouse line is compared to the isogenic control). Graphs show average ± sd, n=3. Statistical differences were determined using a Two-way ANOVA with Tukey’s correction, **p<0.01, ****p<0.0001.

**Fig. S6.**
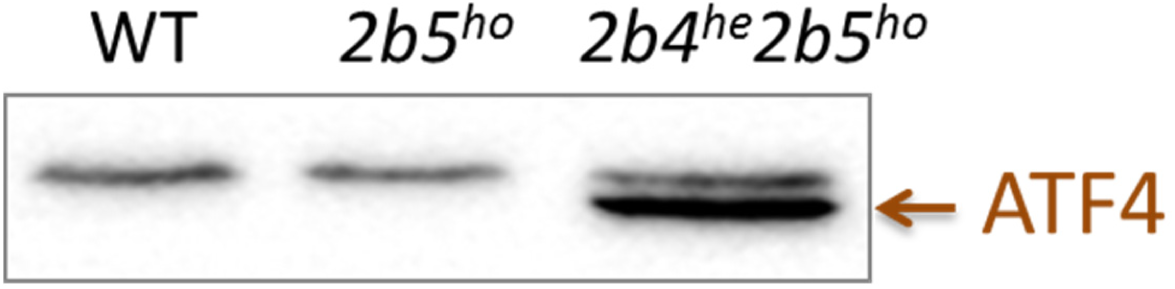
ATF4 protein expression is detected in cerebella nuclei from *2b4*^*he*^*2b5*^*ho*^ mice. Analyses were performed with tissue from 4-month-old mice. Nuclear lysates of mouse brains were obtained by dounce-homogenizing freshly obtained brain tissue in 1 ml PBS with protease inhibitors (PI) and phosphatase inhibitors (PPI), basically as described [23]. The suspension was centrifuged for 5 min at 4000 rpm at 4 °C. The pellets were washed with PBS/PI/PPI and subsequently lysed in extraction buffer (20 mM Tris pH 7.5, 12% glycerol, 125 mM NaCl, 5 mM MgCl2, 0.2 mM EDTA, 10 mM DDT, 0.1% NP40) and sonicated for two brief pulses on ice. The lysates were centrifuged for 10 min at 10,000 xg at 4 °C. Equal amounts of proteins were loaded onto the gel. The upper band is a non-specific protein. ATF4 detection in total tissue samples (nuclei and cytoplasm) showed many non-specific protein signals. The detection in 2b5ho tissue is likely below threshold, as ATF4 was detected with IHC in brain tissue from all VWM genotypes as well as its regulated protein 4E-BP1.

**Fig. S7.**
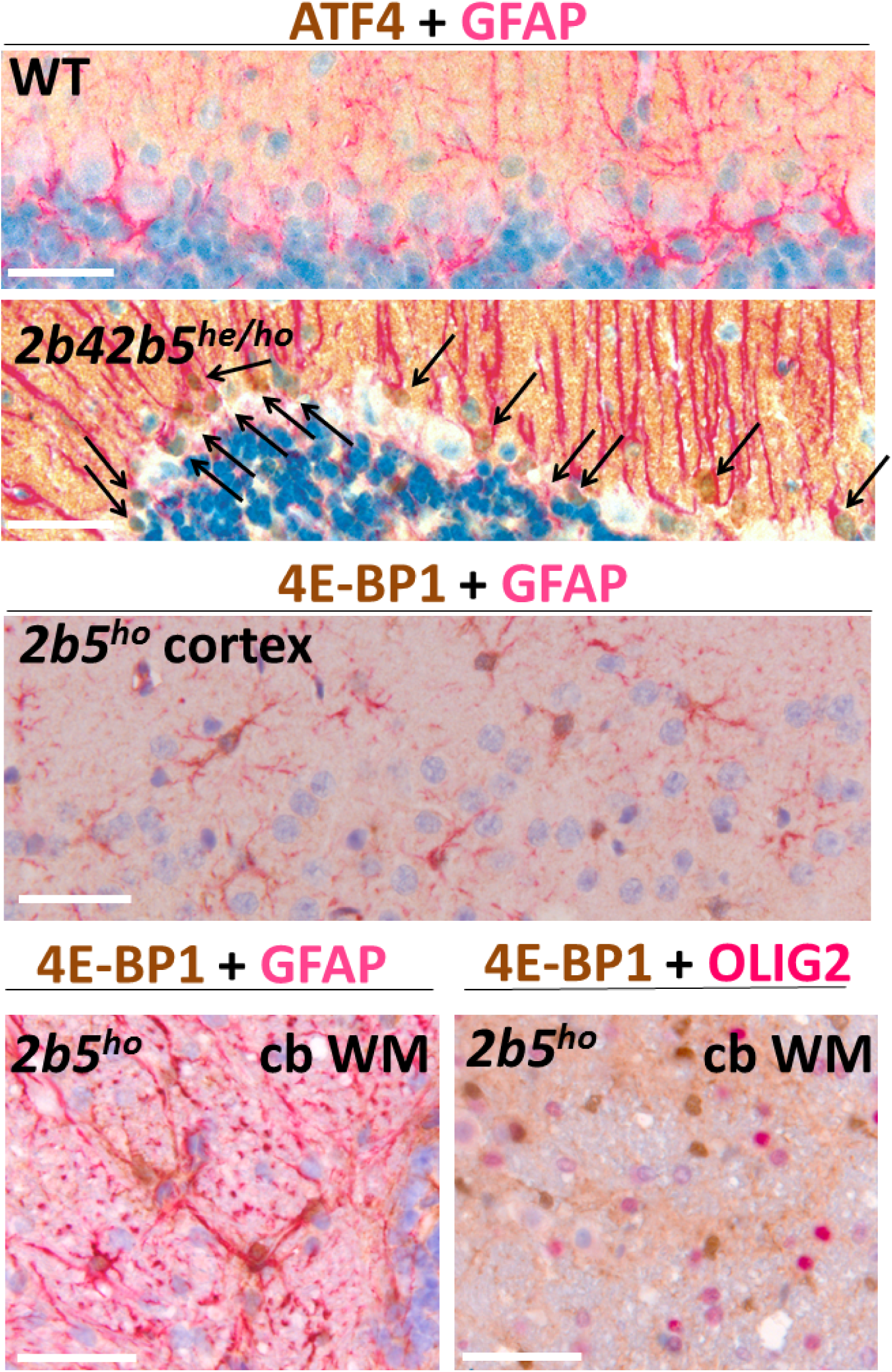
Co-staining for ATF4 or 4E-BP1 with GFAP or OLIG2 demonstrate double positive cells of astrocyte but not oligodendrocyte lineage in VWM mice. ATF4-GFAP co-staining was only successful on sections from *2b42b5*^*he/ho*^ mice (4-month-old). Arrows indicate nuclei of double positive Bergmann glia. 4E-BP1+GFAP staining of astrocytes in cortex is at 2-month-old age. 4E-BP1+GFAP and 4E-BP1+OLIG2 double staining of cerebellar white matter (cb WM) is at an age of 12 months (HEP, *2b5*^*ho*^). Double positive astrocytes were prominent, whereas oligodendrocytes were not detected, even at HEP.

**Fig. S8.**
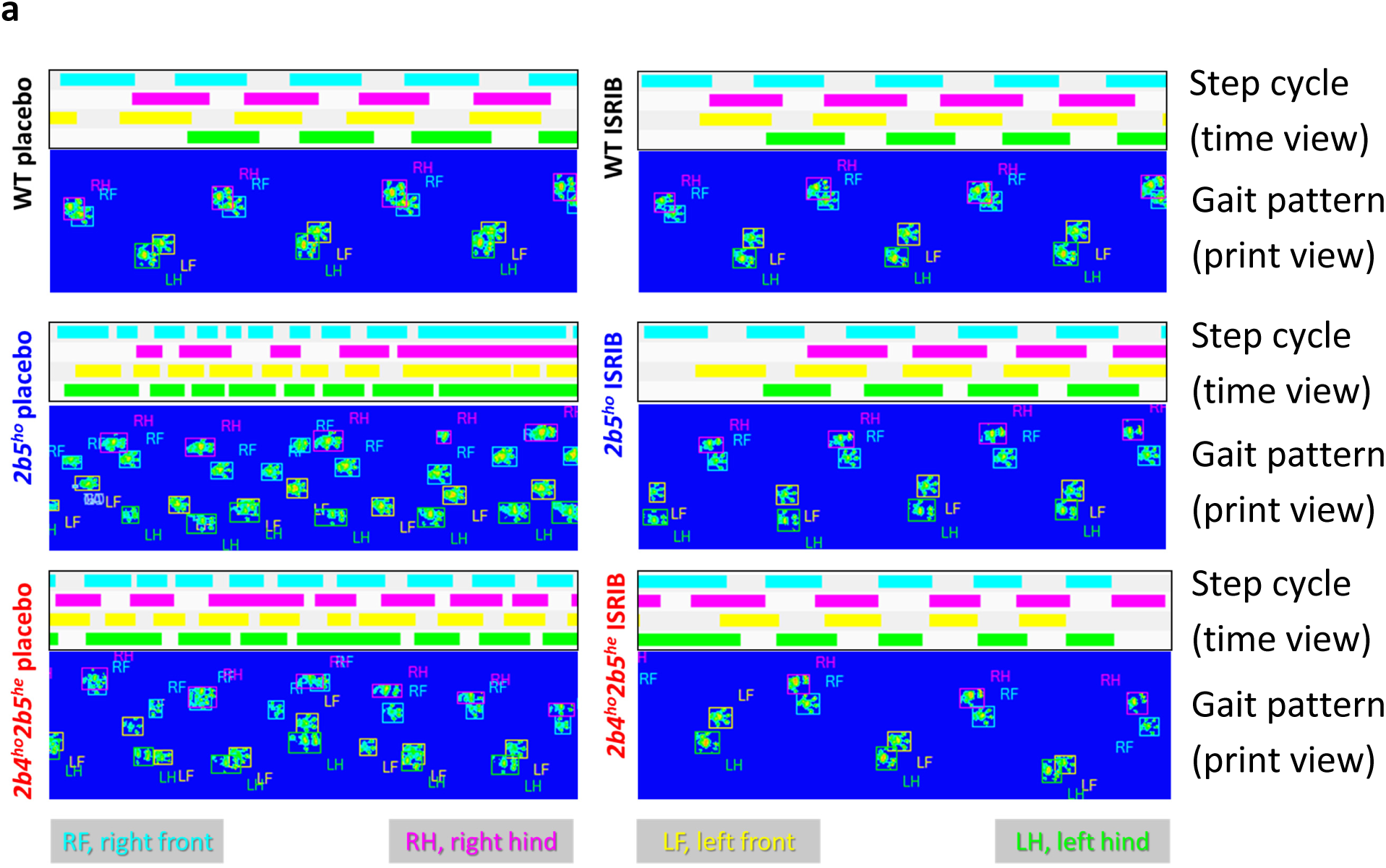
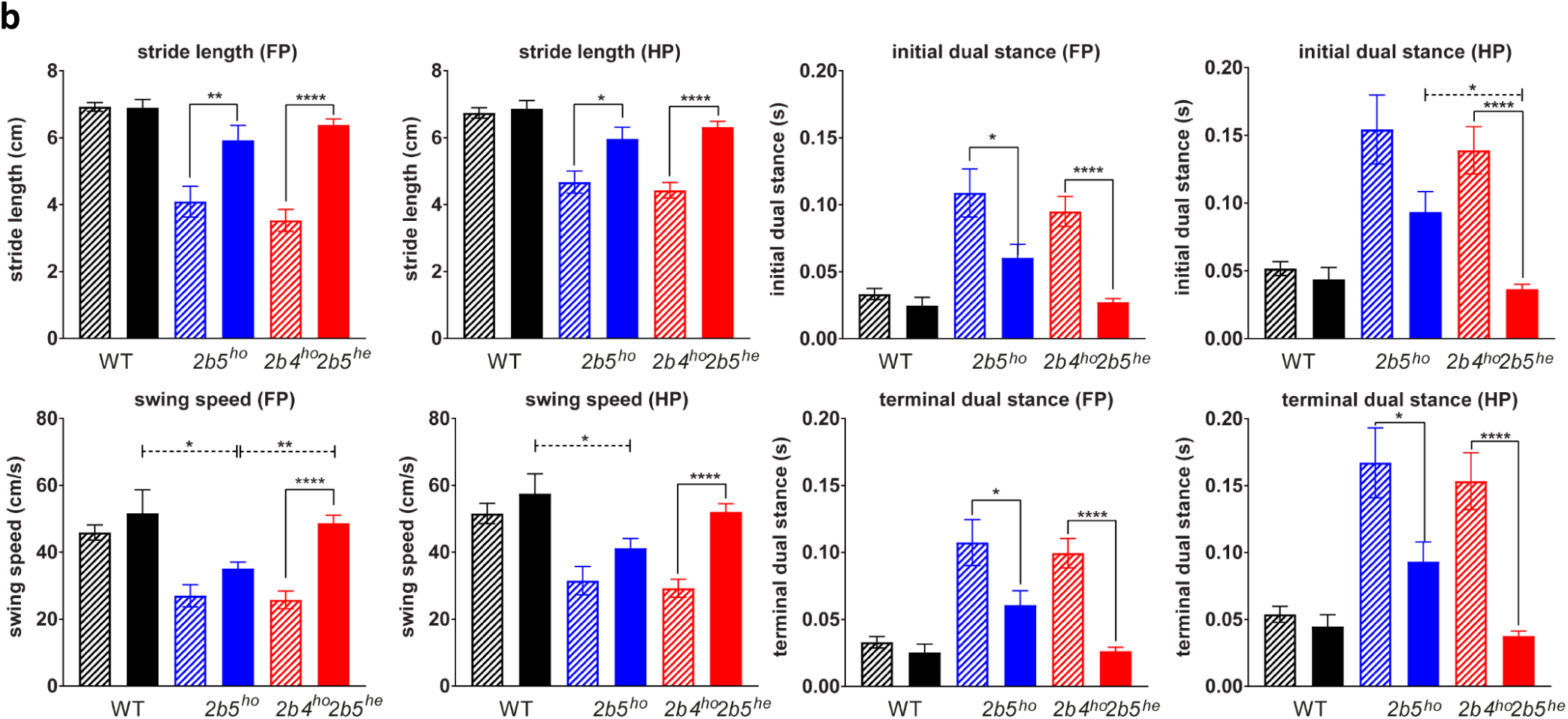
ISRIB ameliorates gait abnormalities in two VWM mouse models, most effectively in *2b4*^*ho*^*2b5*^*he*^ mice; additional data of WT and VWM mice treated with placebo or ISRIB. (**a**) Exemplary data on CatWalk for one mouse within each placebo- and ISRIB-treated group. Representative trace of the step cycle (time view, with colors indicating time that each paw is in contact with the glass plate) combined with the accompanying gait pattern (print view) are indicated. Genotypes and treatments are shown on the left-hand side (WT, black; *2b5*^*ho*^, blue; *2b4*^*ho*^*2b5*^*he*^, red). (**b**) Each paw is color-coded as indicated below the traces. Additional quantified parameters from CatWalk tests not displayed in Figure 4. Error bars indicate SEM. Statistical analysis investigating the ISRIB differences within and among WT, *2b5*^*ho*^ and *2b4*^*ho*^*2b5*^*he*^ was performed with a two-way ANOVA with Tukey’s correction (Supplementary Data File 5). *p<0.05, **p<0.01, ***P<0.001, ****p<0.0001.

**Fig. S9.**
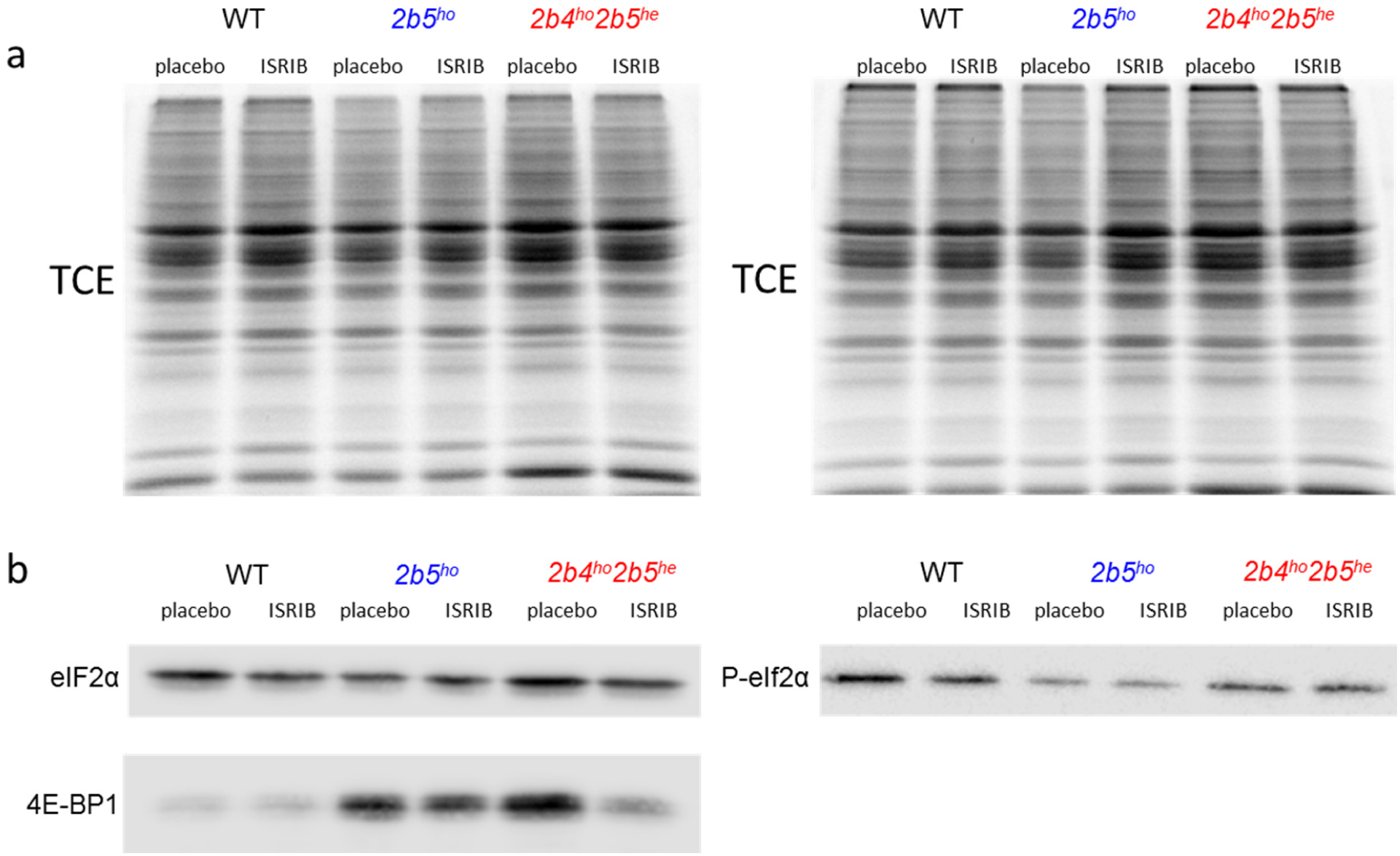
eIF2α phosphorylation is unaltered and 4E-BP1 protein levels are reduced in VWM mouse cerebellum by ISRIB. a) Equal amounts of protein were applied to SDS-PAGE. In-gel protein loading and sample transfer was checked with 2,2,2-trichloroethanol (TCE) as described (*72*). B) Western blot for eIF2α and phosphorylated eIF2α and 4E-BP1. Quantitative results are shown in Fig. 5a.

**Fig. S10.**
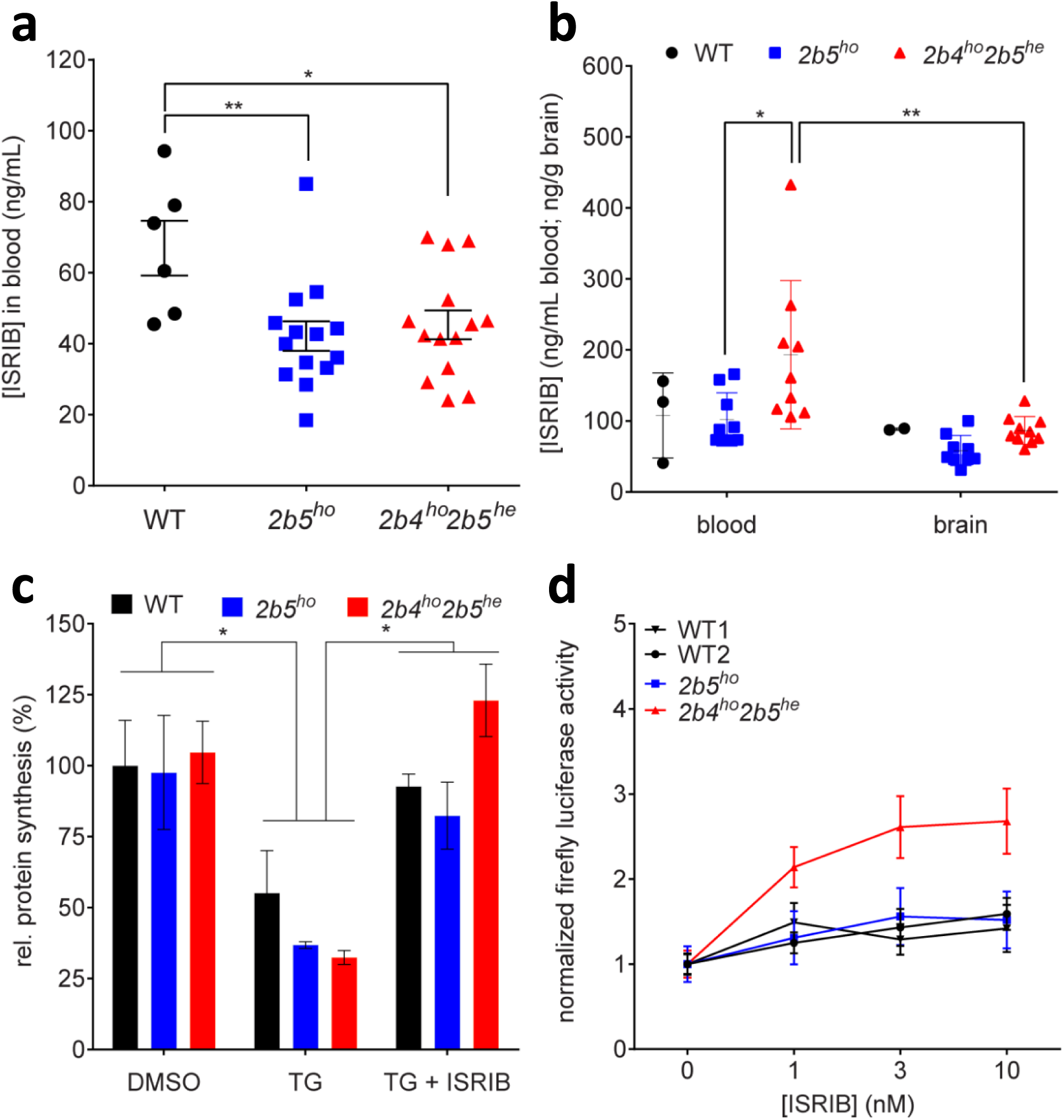
ISRIB’s efficacy in VWM is influenced by the identity of eIF2B mutation. (**a**) ISRIB concentrations were determined in venous blood samples obtained from WT (n=6 per group) and VWM mice (*2b5*^*ho*^ and *2b4*^*ho*^*2b5*^*he*^ n=14 per group) after 14 days of daily ISRIB injections (24 hours after the latest injection); data obtained from placebo-treated animals were below detection and were omitted from the graph. Statistical differences amongst genotypes were determined with One-way ANOVA followed by Tukey’s correction. (**b**) ISRIB concentrations were determined in postmortem blood and olfactory bulb tissue from WT (n=2 per group) and VWM mice (*2b5*^*ho*^ and *2b4*^*ho*^*2b5*^*he*^ n=9 and n=8 per group). Statistical differences were determined with Two-way ANOVA (tissue and genotype as factors) with Tukey’s correction. (**c**) ISRIB restored protein synthesis rates in primary cultures of adult mouse astrocytes undergoing UPR, most effectively in cells from *2b4*^*he*^*2b5*^*ho*^ mice. TG effects were not statistically significant as before (*30*). Statistical differences in test conditions (DMSO vs TG vs TG+ISRIB within WT, *2b5*^*ho*^ and *2b4*^*ho*^*2b5*^*he*^ adult mouse astrocytes were determined with two-way ANOVA with Tukey’s correction (n=2). (**d**) ISRIB enhanced expression of *Gapdh*-driven firefly luciferase reporter in *2b4*^*ho*^*2b5*^*he*^ astrocytes at lower concentrations than in WT or *2b5*^*ho*^ astrocytes (p=0.0026). WT1 and WT2 indicate different isolation dates. Areas under curve (AUC) were calculated for each cell isolate per transfection experiment after correcting for differences between experiments but not between test conditions within experiments (genotype, treatments). Statistical differences between ISRIB sensitivity in WT1, WT2, *2b5*^*ho*^ and *2b4*^*ho*^*2b5*^*he*^ adult mouse astrocytes were determined with one-way ANOVA with Sidak’s correction using AUC values. For all panels, *p<0.05, **p<0.01.

## Supplementary data files

Data file S1: qPCR primers

Data file S2: Polysome association in 2b5ho mouse brain differs for 176 mRNAs (234 probes)

Data file S3: Enrichment of ER stress pathway - Results fishers exact test Mouse and Human

Data file S4: Patient information.

Data file S5: Results statistical analyses.

